# MeCP2 requires interactions with nucleosome linker DNA to read chromatin DNA methylation

**DOI:** 10.1101/2025.06.17.659564

**Authors:** James A. Watson, Beatrice K. Alexander-Howden, Theo S. Hall, Martin A. Wear, Finlay McGhie, Gillian Clifford, Hannah Wapenaar, Juan Zou, Adrian Bird, Marcus D. Wilson

## Abstract

Methyl-CpG-binding protein 2 (MeCP2) is an epigenetic reader essential for neuronal function, but how it binds DNA methylation within chromatin is unclear. Using designer nucleosomes we observe that MeCP2 preferentially engages DNA methylation positioned at multiple sites around the nucleosome. Surprisingly, MeCP2 can bind methylated DNA in bent, histone-contacting core nucleosomal DNA. However, this activity requires additional DNA linker interactions, using MeCP2 regions beyond its canonical methyl binding domain. We mapped a novel DNA-binding region in MeCP2 required for this function. Furthermore, histone H1 antagonises the MeCP2-nucleosome interaction by competing for linker DNA. Overall, this reveals that MeCP2 combines nonspecific but essential interactions with linker DNA to aid specific binding to nucleosomal methylated DNA, independent of nucleosome structure. Our findings reveal a novel contribution to full MeCP2 function, and mechanistic insight into how this clinically important protein interacts with chromatin.

## Introduction

Methyl-CpG-binding protein 2 (MeCP2) is an epigenetic reader of cytosine methylation (meC) which regulates gene expression^1–10^. MeCP2 is highly abundant in neurons^11^ and essential for normal neural maturation and function^12,13^. Indeed, heterozygous mutations cause Rett syndrome^4,14–16^, a severe neurological disorder affecting around 1 in 10,000 live female births^17^. Mouse and cellular models have helped to explain the pathophysiology of Rett syndrome^12,18–20^, but a complete molecular explanation of how this leads to the observed disease phenotypes is lacking.

MeCP2 modulates gene expression and genome architecture via its recruitment to chromatin, and the binding of additional factors^18^. MeCP2 binds chromatin pervasively in the genome through a multifaceted, context-dependent mechanism that is not fully understood^21^. There is evidence that this is guided by the epigenome through DNA methylation recognition^7,11,22–24^, although additional DNA binding specificities have been proposed. MeCP2 contains only a single folded domain, the methyl-binding domain (MBD), that binds DNA cytosine methylation (meC). Upwards of 80% of meCpG sites in the genome are methylated, which is amplified further in neurons with abundant asymmetric meCpA modifications, both of which can be engaged by MeCP2’s MBD^1,5,6,22,24–31^. *In vitro* the highly basic and disordered protein also robustly binds unmethylated double stranded DNA, with preference for methylated DNA^28,32–34^. Outside of the MBD, DNA binding activities have also been identified throughout the rest of the protein, which likely contribute to a general DNA affinity of the protein^1,35–41^.

MeCP2 binding occurs within chromatin, not on naked DNA. The nucleosome is the fundamental unit of chromatin and comprises an octameric core of histones that wrap and compact ∼145-147 bp of DNA, punctuated by linear linker DNA between the repeating nucleosome core particle units^42^. Previous *in vitro* studies of MeCP2 interacting on nucleosomes have proposed that MeCP2 can bind meCpG on solvent-facing nucleosomal major grooves^43,44^. MeCP2 also binds both unmethylated and methylated nucleosomes^44–47^, with different behaviours on chromatin versus linear DNA^48^. Histone proteins, and histone tail modifications, have been suggested as potential targets for MeCP2^45,48–52^. Indeed, in cells MeCP2 binds in a meC-specific and non-specific manner^9,11,23,34^ but the molecular basis for full length MeCP2 interaction with chromatin remains unclear.

We sought to better understand the binding mode of MeCP2 to nucleosomal DNA methylation. Single sites of DNA methylation were engineered throughout the nucleosome, using both meCpG and meCpA sequences. We find that MeCP2 can bind DNA methylation on the nucleosome surface, even when meC sites are facing the histone octamer, which would be predicted to block binding. This unanticipated binding was dependent on the presence of accessible nucleosomal linker DNA and required a central region of MeCP2 outside of the previously characterised domains. Indeed, we find MeCP2 to have little to no binding to nucleosome core particles lacking linker DNA, nor to H1 bound chromatosomes with short linker DNA. Additionally, mutation of this novel DNA binding region disrupted meC specificity *in vitro* and altered MeCP2 binding dynamics to heterochromatin *in vivo*. Our findings suggest that the MBD in isolation, or truncated MeCP2 constructs lacking the central region^53^, would not have access to DNA methylation at all sites in the genome.

## Results

### Full-length MeCP2 is required to bind nucleosomal DNA methylation

We created site specifically-modified designer nucleosomes to probe the interaction of MeCP2 on methylated DNA at specific nucleosome locations. Use of strong positioning DNA sequences^54^ allowed us to precisely situate the histone octamer core with defined lengths of DNA linker regions. We then installed site-specific cytosine CpG DNA methylation (meC), in either the linkers or core-binding DNA (Extended Data Fig. 1A), which could be wrapped into nucleosomes (Extended Data Fig. 1B, Supplementary Fig. 1B-D).

The methylated DNA binding domain (MBD) of MeCP2 is critical to promote interaction with methylated cytosines (meC) in linear DNA (Fig. 1A). We first sought to determine if the MBD in isolation was sufficient to allow meC reading in the context of nucleosomes. Using purified recombinant MeCP2 MBD (residues 77-167, Supplementary Fig. 1A) we tested the interaction by electrophoretic mobility shift assay (EMSA) with nucleosomes containing unequal 37 bp and 27 bp long DNA linker arms and a core sequence of Widom 601 DNA (termed 37-N_601_-27). An unlabelled double stranded DNA competitor was included in all assays to limit non-specific binding for methylation specificity (Supplementary Fig. 1E), as described previously^44,45^.

**Figure 1:**
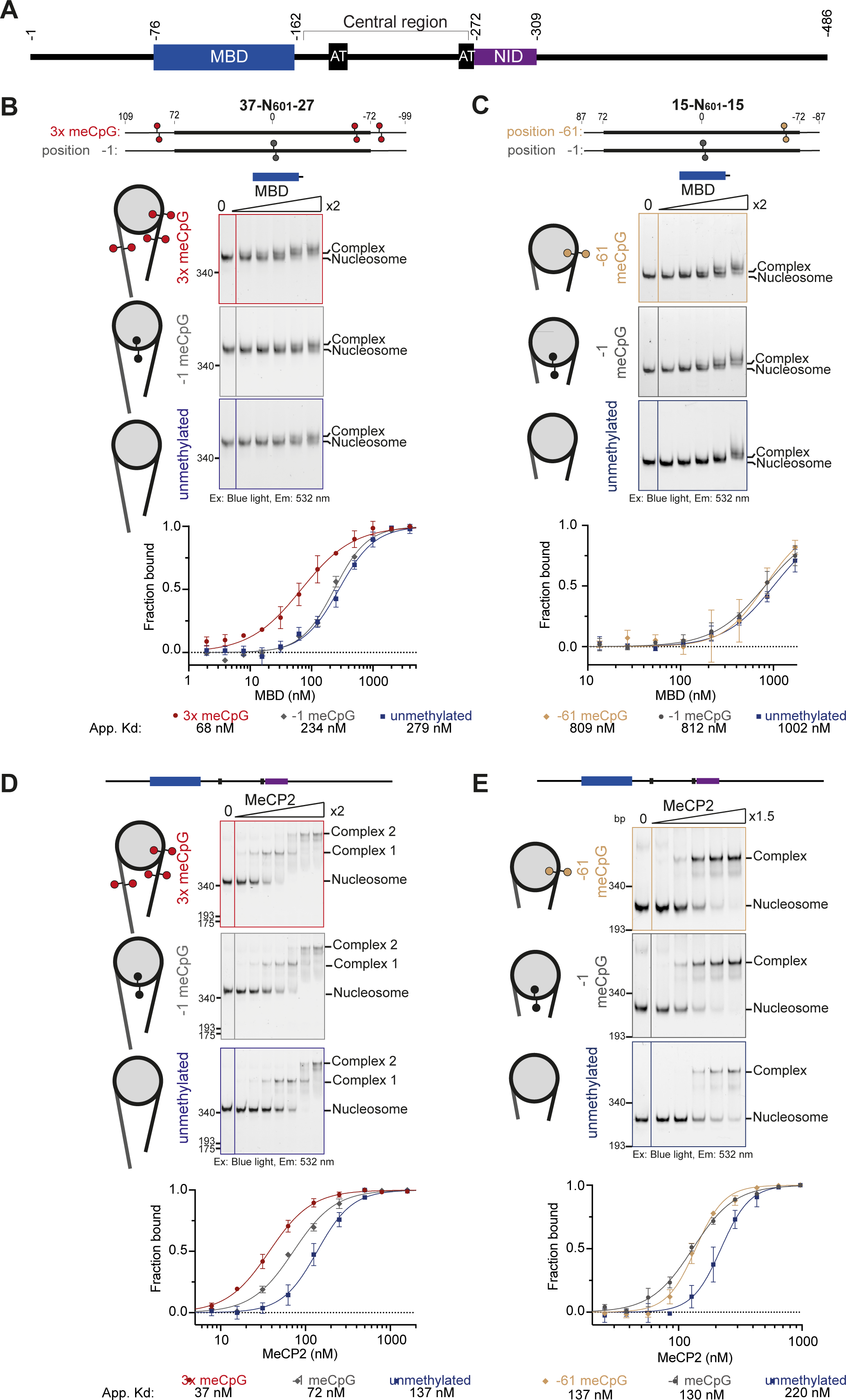
Full-length MeCP2 is required to bind nucleosomal DNA methylation. **A.** Diagram of full-length human e2 MeCP2 highlighting the key methyl binding domain (MBD) (blue box) and NCoR/SMRT interaction domain (NID) (purple box). Domain boundaries are indicated with residue number. AT hook motif (AT) 1 and 2 are also shown (black boxes). **B.** Representative EMSA native gels (3 repeats) showing a 2-fold dilution series of MBD with 37-N_601_-27 nucleosomes (15.6-250 nM). Nucleosomes were either methylated with meCpG at three positions (+83, −61, −80 bp from the dyad) (red), a single meCpG at (−1 bp from the dyad) (grey), or unmethylated (blue). Free nucleosome and complex bands are indicated. Quantification of the free nucleosome bands at each concentration, of the full concentration series, was fitted with a binding isotherm and an apparent dissociation constant (K_D app_) calculated. **C.** Representative EMSA native gels (3 repeats) showing a 2-fold dilution series of MBD interacting with 15-N_601_-15 nucleosomes (107-1713 nM). Nucleosomes were methylated with a single meCpG either −61 bp (brown) or −1 bp (grey) from the dyad, or unmethylated (blue). Binding isotherms and K_D app_ are also shown. **D.** Representative EMSA native gels (3 repeats) showing a 2-fold dilution series of full-length MeCP2 with 37-N_601_-27 nucleosomes (15.6-1000 nM). Binding isotherms and K_D app_ are also shown. **E.** Representative EMSA native gels (3 repeats) showing a 1.5-fold dilution series of full-length MeCP2 on 15-N_601_-15 nucleosomes (85-430 nM). Binding isotherms and K_D app_ are also shown.

As expected, the MBD robustly bound to nucleosomes containing multiple meC sites in the linear linker DNA projecting away from the nucleosome (−80, −61 and 83 bp from the centre point of the nucleosome or dyad; Fig. 1B), with reduced binding to non-methylated nucleosomes. However, placing a single meC site in the extensively bent, histone-contacting nucleosome core (−1 bp from the dyad; Supplementary Fig. 1C & D), no longer allowed methyl-specific binding of the MBD (Fig. 1B). Similarly, shorter-linker nucleosome substrates (15-N_601_-15; Supplementary Fig. 1C & D) also lacked methylation preference when a meC was placed near the dyad (Fig. 1C). This phenomenon was not due to the exact positioning of meC, as moving the meC closer to the entry/exit DNA site of the nucleosome core (position −61, Fig. 1C) also blocked binding. Nucleosomes wrapped with a different sequence (16-N_603_-30; Extended Data Fig. 2A) similarly showed greatly reduced binding affinity for the MBD when meC was positioned closer to the nucleosome core, indicating that the effect was independent of local-sequence context. We conclude that methylated DNA sites associated with the nucleosome core are refractory to meC binding by the MBD.

In contrast to the MBD alone, full-length MeCP2 was indifferent to meC positioning on a nucleosome. Specifically, purified MeCP2 (Supplementary Fig. 1A) could preferentially recognise DNA methylation on linker DNA but surprisingly also within the histone-bound nucleosome core DNA (Fig. 1D & E). Likewise, unlike the MBD, MeCP2 did not display reduced binding to linker meC close to the nucleosome-core (Extended Data Fig. 2A). Swapping asymmetric meCpA sites for meCpG yielded similar DNA methylation specificity (Extended Data Fig. 2B). Using full length MeCP2 increased overall affinity for meC nucleosomes, suggesting additional binding capability. However, MeCP2 also reduced the selectivity to meC compared to the MBD, suggesting this added affinity is not DNA methylation sensitive. Interestingly, MeCP2 displayed a generally stronger binding affinity to nucleosomes containing CpA sequences over CpG, independent of methylation, suggesting a potential sequence preference^34^.

In all cases MeCP2-nucleosome complexes migrated as distinct individual bands. On longer nucleosome substrates a second slower migrating band was also observed at high MeCP2 concentrations, suggesting a second binding event could be accommodated by increased DNA linker length. Mass photometry analysis of complexes confirms the presence of two MeCP2 binding events to 37-N_601_-27 nucleosomes, and a single binding event to 15-N_601_-15 nucleosomes (Extended Data Fig. 2C). Taken together, our results suggest that meC recognition within nucleosome cores by MeCP2 requires regions in addition to the MBD.

### MeCP2 requires linker DNA to bind meC sites in core nucleosomal DNA

Full length MeCP2 displayed higher binding affinity to both meC- and non-methylated nucleosomes compared to the MBD alone. We first tested if this was due to histone protein interactions^45,48–52^. Crosslinking mass spectrometry shows links between the MBD and N-terminal tail of H3 (Extended Data Fig. 3A). However, the H3 tail is adjacent to the engineered sites of meC, and shifting DNA methylation into the core of the nucleosome also varied the protein-protein crosslinking pattern, reflecting the location of meC rather than a consistent feature of MeCP2 binding. Furthermore, removing the tail of H3 or addition of H3K_c_27me3 - which has been previously suggested to be a binding target of the MBD^50,51^, as well as antagonise MeCP2-histone binding^52^ - did not affect MBD binding to nucleosomes in these assays (Extended Data Fig. 3B). Indeed, removal of the H3 tail globally increased MeCP2 binding to nucleosomes (Extended Data Fig. 3C). The H3 tail is known to bind linker DNA^55,56^, raising the possibility that the histone tail competes with a MeCP2-linker binding that is important for MeCP2 interaction with the nucleosome.

To test the hypothesis that MeCP2 binding to DNA on the nucleosome surface requires linker DNA, we assayed the MeCP2 interaction with linker-less nucleosomes. Strikingly, we found that removal of linker DNA to form nucleosome core particles (N_601_) disrupted binding of MeCP2, with or without meC (Fig. 2B). This linker requirement was independent of DNA methylation. As longer linker lengths enhanced binding of full length MeCP2 (Fig. 1D & E), we therefore hypothesised that additional DNA binding aids MBD-meC chromatin interaction. MeCP2 contains appreciable non-specific DNA binding ability which drives dynamic association with the genome^39,57^. Indeed, direct comparison of MeCP2 binding to DNA of various lengths, either as wrapped nucleosomes or unwrapped free DNA, highlights its binding preference for linear double stranded DNA (Extended Data Fig. 4A).

**Figure 2:**
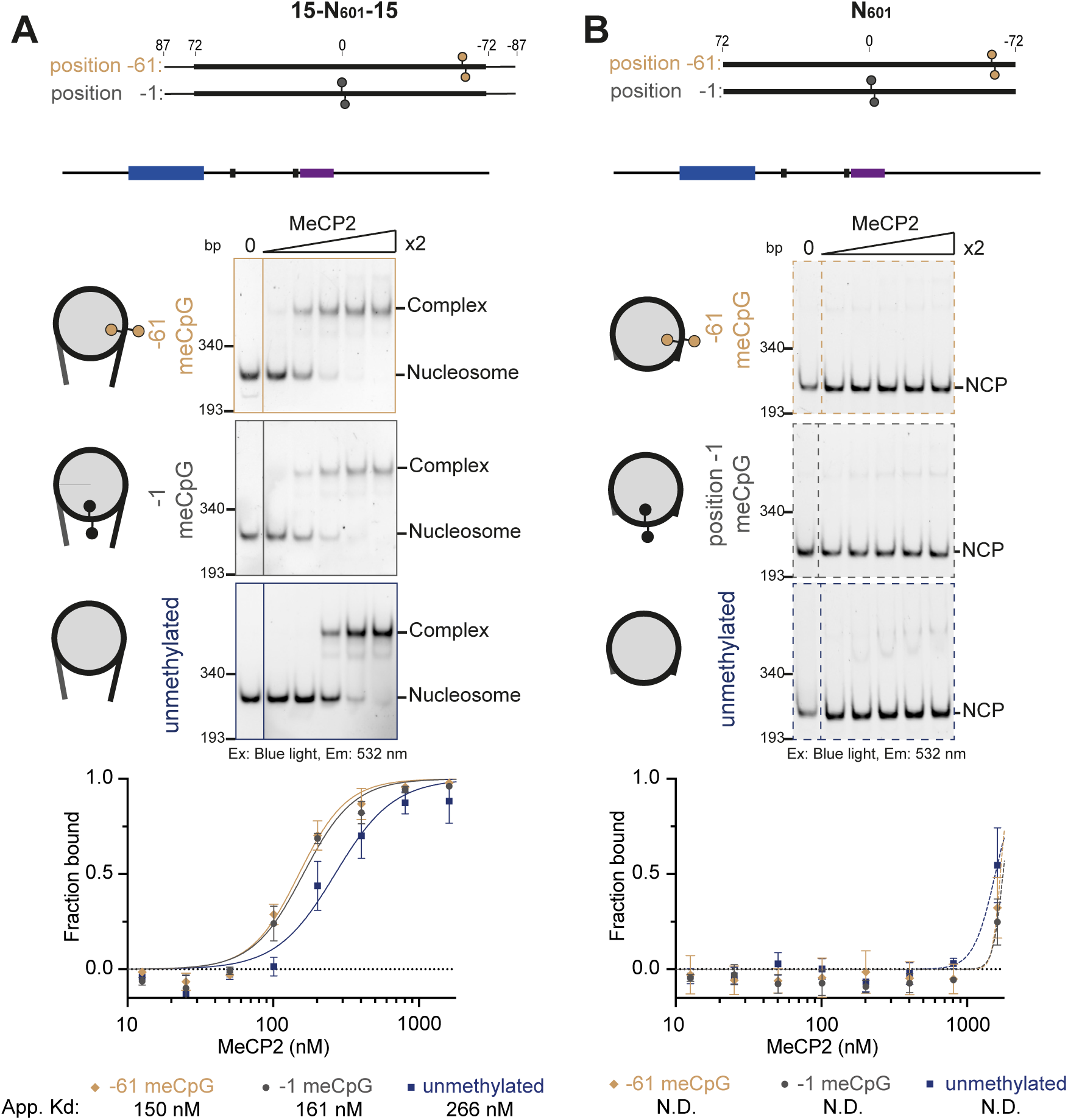
Linker DNA is essential for MeCP2 to bind nucleosomal DNA methylation. Representative EMSA native gels (3 repeats) showing a two-fold dilution series (50.4-807 nM) of MeCP2 on 15-N_601_-15 nucleosomes (**A**) and N_601_ nucleosome core particles (**B**). Nucleosomes were methylated with meCpG either at position −1 (grey) or −61 (brown), or unmethylated (blue). Free nucleosome, nucleosome core particle (NCP) and complex bands are indicated. Binding isotherms and K_D app_ are also shown.

To further examine the importance of nucleosome linker DNA to MeCP2 binding, asymmetric nucleosomes with only a single linker were tested for MeCP2 interaction. The presence of a single 15 bp linker (N_601_-15) led to overall poorer binding and a loss of meC specificity compared to 15-N_601_-15 (Extended Data Fig. 4B). Extending the linker (N_603_-30), retained both DNA binding and meC specificity and produced similar binding profiles compared to 16-N_603_-30 (Extended Data Fig. 4C), suggesting the single 30 bp linker was sufficient to maintain optimal MeCP2 binding to nucleosomes accommodating both the MBD and DNA interacting region of MeCP2 (Extended Data Fig. 4D). This suggests that the MBD of full length MeCP2 can no longer engage robustly without the stability conferred by additional DNA binding elements and MeCP2 combines meC-MBD and DNA linker interaction.

### A central region of MeCP2 binds nucleosome linker DNA

The data above suggests that DNA binding capability outside of the MBD is required for MeCP2 to gain access to meC in a nucleosome core. To test this we next set out to identify which regions of MeCP2 are involved in nucleosome linker binding. Crosslinking mass spectrometry suggested a central region of MeCP2 is adjacent to the nucleosome (Extended Data Fig. 3A). However, the histone acidic patch, a common site of chromatin-protein engagement^58^, was not required for the interaction in this context (Extended Data Fig. 5A-C). The central region is sometimes referred to as the Intervening Domain (ID) and Transcriptional Repression Domain (TRD), and contains reported DNA binding activity at AT-hooks 1 and 2^1,35–37,39–41,59^. A construct covering this central region (residues 162-309) bound poorly to nucleosomes core particles (N_601_), but better to nucleosomes with linkers (15-N_601_-15, Fig. 3A & B, Extended Data Fig. 5A & B). Comparing the binding of this central region construct with that of MeCP2 or the MBD alone revealed that the full-length MeCP2 interaction could be explained by a combination of both meC-specific and linker DNA-binding modalities. While full length MeCP2 preferentially binds methylated DNA it shows appreciable affinity for unmethylated nucleosomes (Fig. 3A, Extended Data Fig. 5D). Methylation preference is driven by the MBD: incorporation of a mutation known to ablate meC recognition (R133G) removes discrimination between meC and unmethylated nucleosomes and the MBD alone has very low affinity to unmethylated nucleosomes. Increasing the length of nucleosome linker DNA improves MeCP2 interaction. Notably, the central region shows no methylation-dependant binding preference, but has a clear increased affinity for longer linkers (Fig. 3A & B). The data are compatible with the view that this region engages linker DNA and combines with the MBD to promote meC-nucleosome recognition.

**Figure 3:**
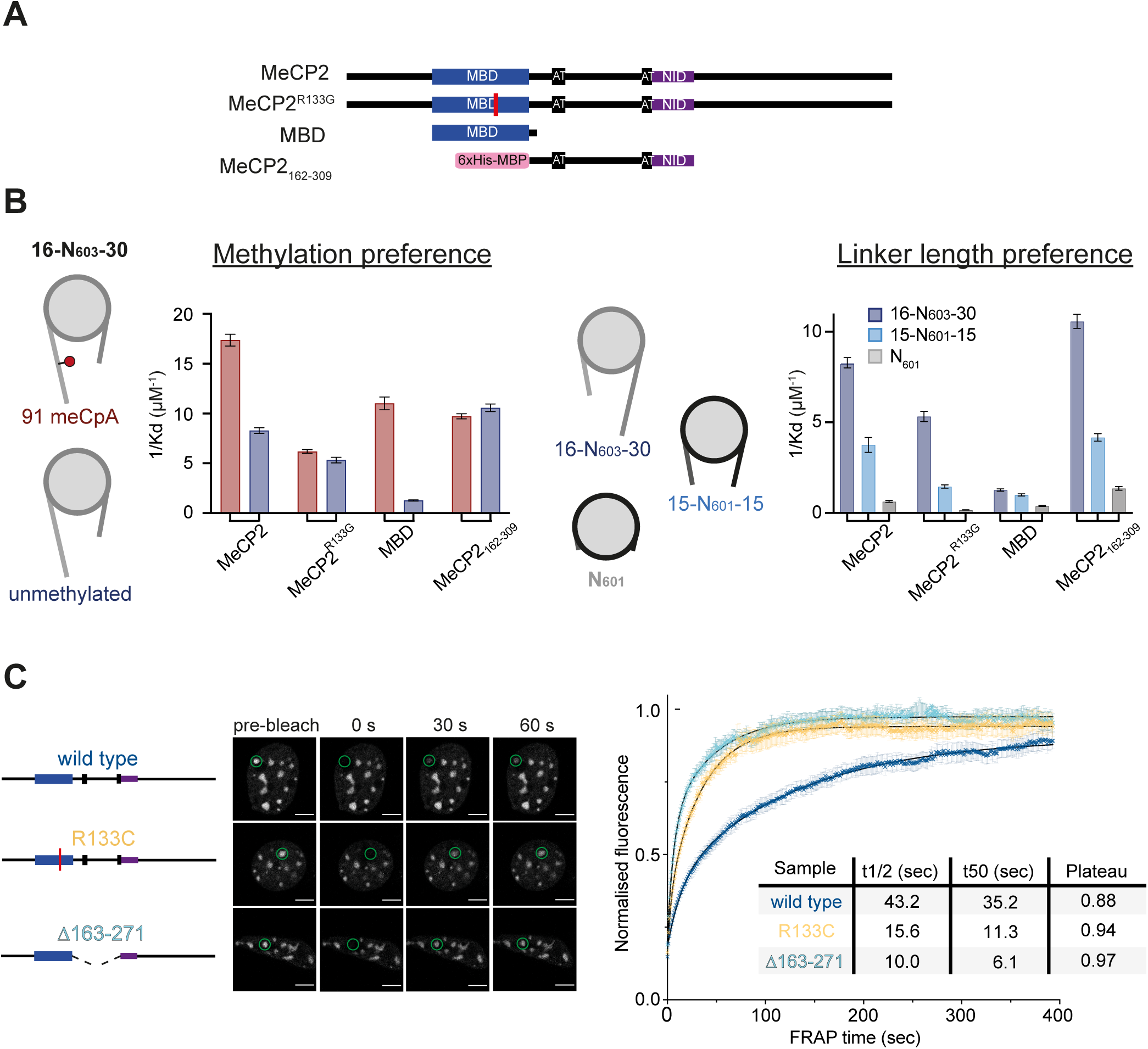
A central region of MeCP2 binds nucleosome linker DNA. **A.** Diagram of MeCP2 constructs used in part **B**. The domains of MeCP2, and tags, present in each construct is highlighted. MBD mutation R133G location is marked by a red line. **B.** Summary of MeCP2 variant affinity measurements determined by EMSA. (Left) Methylation preference was assayed by comparison of binding affinities for each construct on 16-N_603_-30 nucleosomes, with either 91 meCpA or unmethylated. (Right) Linker length preference by comparison of affinities on unmethylated 16-N_603_-30, 15-N_601_-15 and N_601_ nucleosomes. The summary of each data set is shown by plotting the inverse apparent dissociation constants (1/Kd). **C.** FRAP quantification of wild-type (blue), Δ163-271 (cyan) and R133C (yellow) eGFP-MeCP2 recovery in mouse fibroblasts. The number of analysed cells, from 3 independent experiments, are: WT *n = 37* cells, Δ163-271 *n* = 33 cells, R133C *n* = 28. Error bars show SEM. An example time-course from the live cell imaging of each construct is also shown with the bleached foci circled.

To test the importance of the central region for MeCP2 function we performed fluorescence recovery after photobleaching (FRAP) experiments in NIH3T3 mouse fibroblast cells (Fig. 3C, Extended Data Fig. 8). As seen in previous studies wild-type eGFP tagged MeCP2 localised to highly methylated pericentric heterochromatic foci and after bleaching undergoes slow and incomplete recovery, suggesting a stably bound fraction^22,59–62^. Deletion of MeCP2 163-271, covering the central region, resulted in a more complete and rapid recovery, similar to a known Rett syndrome mutation in the MBD (R133C). This suggests that this region is needed for MeCP2 to stably associate with chromatin *in vivo*.

### A novel DNA interacting motif contributes to linker DNA binding activity in central MeCP2

Primary sequence analysis highlighting pathogenic and likely-pathogenic point mutations^63^, sequence conservation^64^, and predicted pathogenicity using AlphaMissense^65,66^, reveals the crucial role of the MBD and NCoR/SMRT interacting domain (NID, Fig. 4A). In addition, the central region between the MBD and NID domains contains several neurodevelopmental-implicated patient mutations corresponding to the AT-hooks 1 and 2, as well as a highly-conserved and invariant uncharacterised region between residues 205-257^16,63,67,68^.

**Figure 4:**
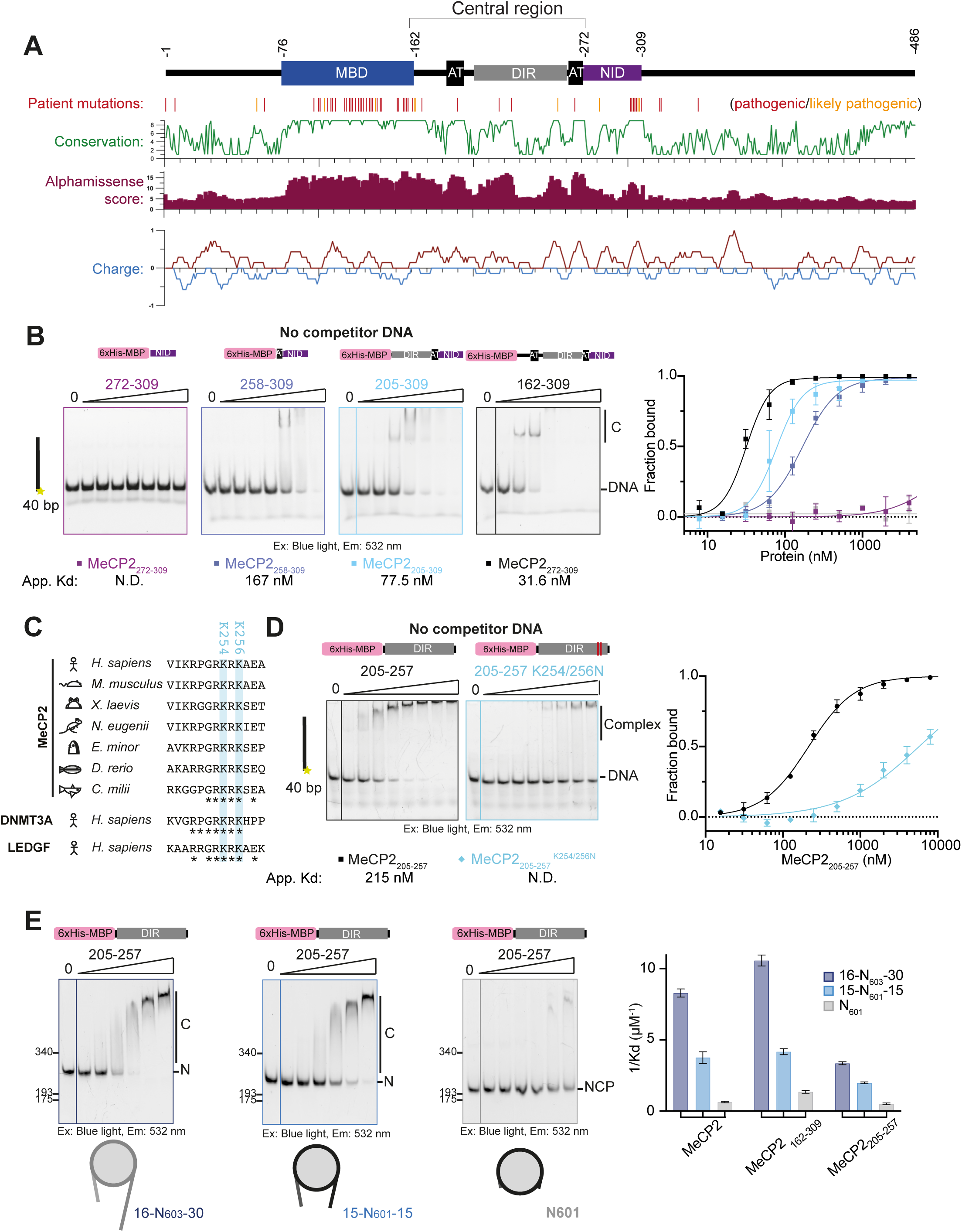
Linker DNA binding occurs through multiple motifs in central region MeCP2, including a novel DNA interacting region. **A.** Schematic summarising bioinformatic analysis of MeCP2. Pathogenic (red), and likely pathogenic (orange), point mutations (Clinvar) plotted along the length of MeCP2. Sequence conservation scores (green), summed AlphaMissense pathogenicity scores (purple), and charge (red-blue) were also plotted. **B.** Representative EMSA native gels (3 repeats) showing a two-fold dilution series (15.6-1000 nM) of each HisMBP tagged MeCP2 construct incubated with 40 bp dsDNA. Free DNA and complex bands are indicated. Binding isotherms and K_D app_ are also shown. No competitor DNA was used in this assay. **C.** Multiple sequence alignments of MeCP2_250-256_, from the DNA interacting region of *Homo sapiens* MeCP2, compared to a variety of animal species. The two lysine residues mutated in experiments are highlighted in blue. **D.** Representative EMSA native gels (3 repeats) showing a two-fold dilution series (31.3-4000 nM) of HisMBP-MeCP2_205-257_ constructs incubated with 40 bp DNA. Free DNA and complex bands are indicated. Binding isotherms and K_D app_ are also shown. No competitor DNA was used in this assay. **E.** Representative EMSA native gels (3 repeats) showing a two-fold dilution series (62.5-2000 nM) of HisMBP-MeCP2_205-257_ on unmethylated 16-N_603_-30, 15-N_601_-15 and N_601_ nucleosomes. The Inverse apparent dissociation constants (1/Kd) is plotted as in (Fig. 3A). Full-length MeCP2 and HisMBP-MeCP2_162-309_ are shown again for comparison.

Using this information, we tested the DNA binding capacity of the central region of MeCP2 using a series of additive purified central region constructs (Supplementary Fig. 2C) in EMSA assays (Fig. 4B). MeCP2_272-309_, covering only the NID of MeCP2, did not bind to DNA at the concentrations used, similar to a His-MBP tag control (Fig. 4B, Extended Data Fig. 6A). Addition of the region containing AT-hook 2 (MeCP2_258-309_) induced DNA binding activity. Extending a further 53 amino acids N-terminal to include the newly identified region (MeCP2_205-309_), further improved binding. This suggests that this region - hereafter termed the DNA Interacting Region, DIR - also contributes to MeCP2 DNA binding activity. Finally, extension to include AT-hook 1 (MeCP2_162-309_) showed the most robust DNA binding activity, suggesting three central MeCP2 DNA-binding regions work together to engage DNA.

A small construct containing only the DIR (MeCP2_205-257_, Supplementary Fig. 2D) was sufficient to bind double-stranded DNA (Fig. 4D), with a preference for longer lengths of DNA (Extended Data Fig. 6B) and no observable DNA sequence preference (Extended Data Fig. 6C). The DIR fragment appears as a monomer in isolation and likely exists as an extended non-globular structure (Extended Data Fig. 6D). The DIR is not overall strongly positively charged (Fig. 4A) but includes a conserved motif ‘R-P-G-R-K-R-K’ (residues 250-256, Fig. 4C). Mutation of two lysine residues (K254N, K256N) significantly reduced DNA binding (Fig. 4D), suggesting that this motif is principally responsible for DNA binding activity of the DIR. Interestingly an identical short amino acid sequence is found in the *de novo* methyltransferase DNMT3A, and a similar motif is also present in Lens Epithelium-Derived Growth Factor (LEDGF/p75) (Fig. 4C). In both cases the regions have been implicated in DNA binding^69–71^. Indeed, we observed that mutation of the motif in a DNMT3A construct also reduced DNA binding (Extended Data Fig. 6E). Both DNMT3A^72^ and LEDGF^73^ also engage with nucleosomes, raising the possibility that this motif could be a generalisable DNA binder that promotes multivalent chromatin interactions. Compatible with this notion, we found that the DIR also bound to nucleosomes with a preference for longer linker DNA, albeit weaker than the entire central region (Fig. 4E). Interestingly, the DIR bound nucleosome core particles poorly, suggesting it does not engage with core nucleosomal DNA, or the histone acidic patch. Therefore, the behaviour of the DIR engaging linear accessible linker DNA mimics the specificity of intact MeCP2.

### Motifs in central MeCP2 aid MBD binding to nucleosomal DNA methylation

If the DNA binding activity of central MeCP2 is necessary to allow binding to meC nucleosomes, we hypothesised that disruption of this region would produce a protein that, like MBD alone, would not be able to access nucleosome core DNA meC. To test this, we purified full-length MeCP2 containing mutations in the DIR region (K254N, K256N) in addition to both AT-hook 1 (R188G, R190G) and 2 (R268Q)^39,60^, to abrogate DNA binding activity in the region (MeCP2 AT-DIR^mut^; Supplementary Fig. 2E). Binding to methylated nucleosomes was assayed *in vitro* as before by EMSA, as well as by both surface plasmon resonance (SPR) and microscale thermophoresis (MST). Reduced DNA binding capabilities of MeCP2 AT-DIR^mut^ ablated overall binding affinity to all 37-N_601_-27 nucleosomes, as expected (Fig. 5A, Extended Data Fig. 7A). Additionally, the mutations had an even greater effect when a single meC was positioned in the nucleosome core: meC preference was weaker compared to the retained activity when meC was in distal linker regions. A similar loss of nucleosomal methylation preference for MeCP2 AT-DIR^mut^ was also seen for binding to 15-N_601_-15 nucleosomes with meC confined to the nucleosome core (Fig. 5B, Extended Data Fig. 7B, Supplementary Fig. 3), as well as to 16-N_603_-30 nucleosomes (Extended Data Fig. 7C). Thus the specificity of full-length MeCP2 with this combination of central region mutations resembles that of the MBD alone, supporting the hypothesis that the central region is responsible for providing access to nucleosomal DNA.

**Figure 5:**
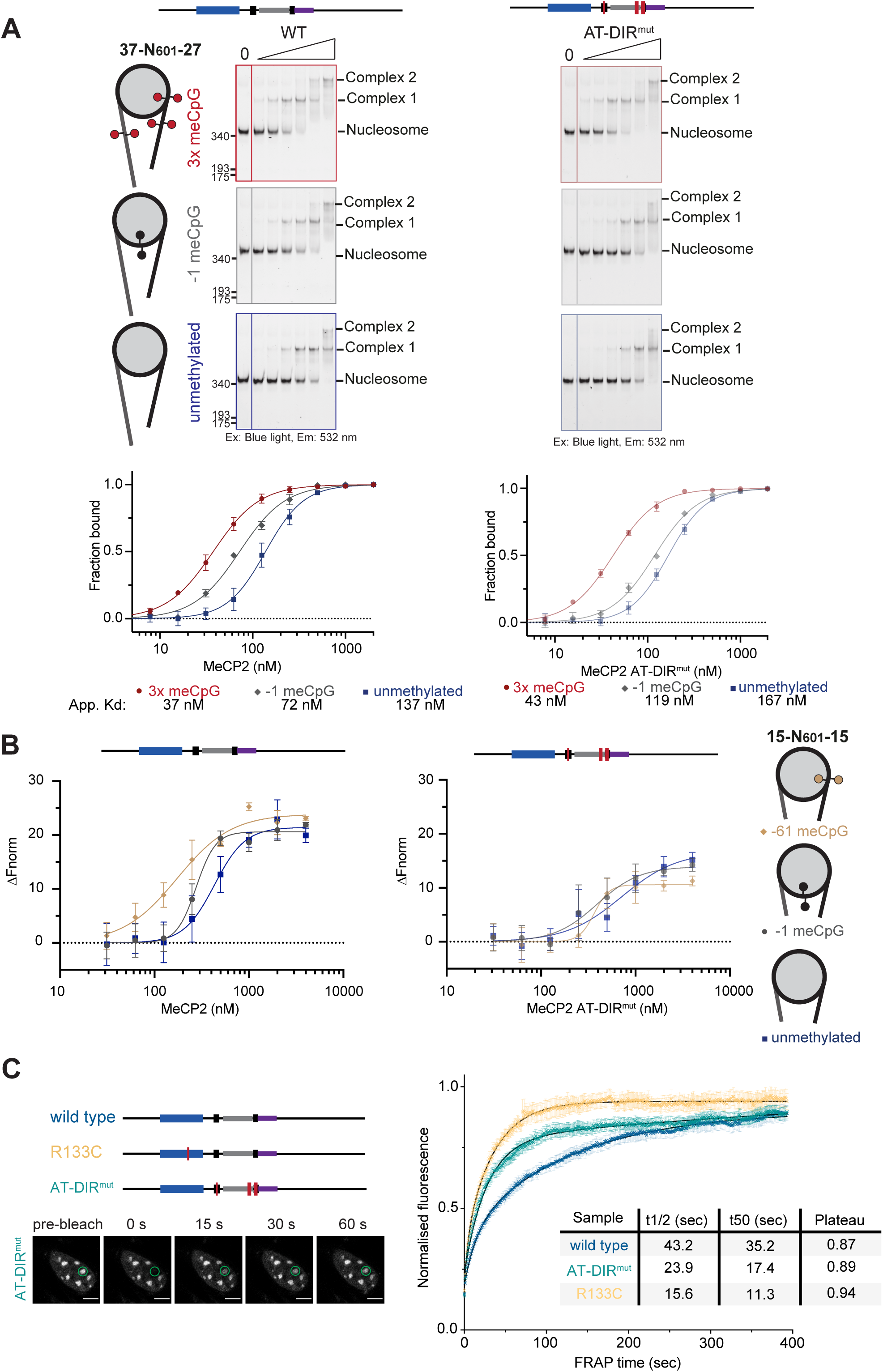
Disruption of DNA interacting motifs in central MeCP2 limit binding to nucleosomal DNA methylation. **A.** Representative EMSA native gels (3 repeats) showing a 2-fold dilution series (15.6-500 nM) of MeCP2 wild-type (left) or AT-DIR^mut^ (R188G, R190G, K254N, K256N, R268Q) (right) with 37-N_601_-27 nucleosomes. WT data was shown previously (Fig. 1D). Nucleosomes were methylated with meCpG at three positions (+83, −61, −80 bp from the dyad) (red), a single meCpG (−1 bp from the dyad) (grey), or unmethylated (blue). Binding isotherms and K_D app_ are also shown. **B.** Amplitude normalised and baseline subtracted MST data (3 repeats) showing a 2-fold dilution series (31.3-4000 nM) of MeCP2 wild-type (left) or AT-DIR^mut^ (R188G, R190G, K254N, K256N, R268Q) (right) on 15-N_601_-15 nucleosomes. Nucleosomes were methylated with a single meCpG either at position −61 bp (brown) or −1 bp (grey), or unmethylated (blue). **C.** Graph showing the FRAP quantification of wild-type (blue), AT-DIR^mut^ (teal) and R133C (yellow) eGFP-MeCP2 in mouse fibroblasts. Wild-type and R133C are shown as before (Fig.3.B.) for comparison. The number of analysed cells, from 3 independent experiments, are: AT-DIR^mut^ *n* = 31 cells. Error bars show SEM.

FRAP analysis of the AT-DIR^mut^ construct in live cells confirmed disrupted chromatin binding dynamics. The magnitude of the effect was similar to, although somewhat milder than, the previously observed MeCP2 central deletion (Δ163-271) and the Rett syndrome causing disease mutation control (R133C; Fig. 5C, Extended Data Fig. 8). In all mutant cases the recovery was more rapid and complete than the wild-type, indicating reduced stability of the MeCP2-chromatin interaction *in vivo*.

### The C-terminal tail of H1 blocks MeCP2 access to nucleosome linker DNA

Linker histone H1 also binds nucleosome linker DNA and is found at a high concentration in neurons, approximately equivalently to that of MeCP2. The two proteins have been reported to condense chromatin and compete with one another for linker DNA binding, although this has been contested^11,44,48,74^. We decided to test whether H1 affects MeCP2 binding to nucleosomes using our designed system. A set concentration of linker H1.0, which is the predominant variant in neurons^75,76^, was first incubated with 15-N_601_-15 nucleosomes to form chromatosomes (Extended Data Fig. 9A & B). No binding of MeCP2 to 15-N_601_-15 chromatosomes was observed even in the presence of nucleosome core meC (Fig. 6A), reminiscent of the absence of binding seen to N_601_ nucleosome core particles. Removal of the C-terminal tail of H1.0, which binds linker DNA, mostly restored MeCP2 binding to chromatosomes (Fig. 6B, Extended Data Fig. 9C). This suggest that H1 can block meC reading in a nucleosome by masking short DNA linkers from MeCP2.

**Figure 6:**
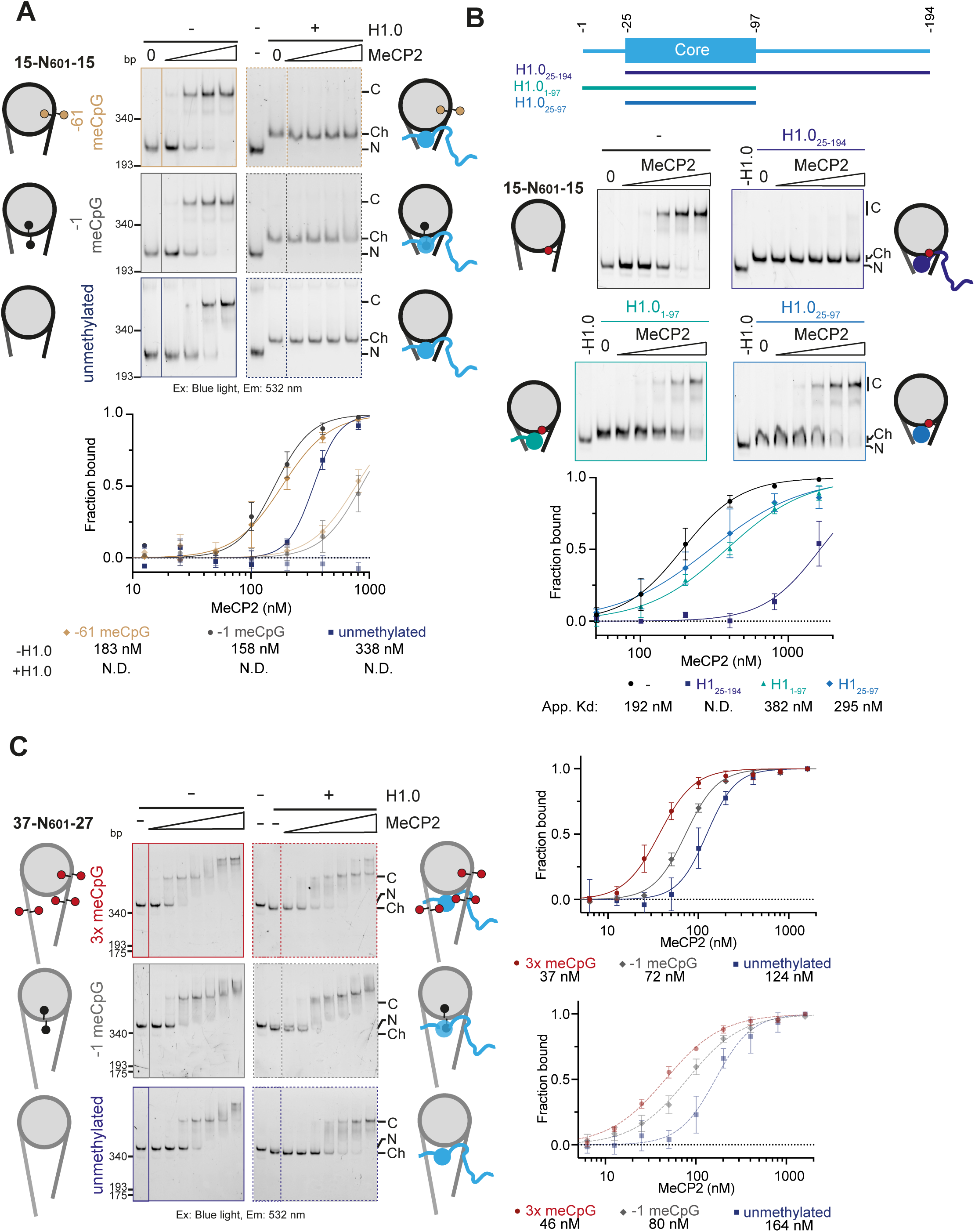
MeCP2 competes with H1.0 for nucleosome linker DNA binding. **A.** EMSA native gels showing a two-fold dilution series (50.4-404 nM) of MeCP2 on 15-N_601_-15 nucleosomes (left) or chromatosomes (right) (3 repeats). Free nucleosome (N), chromatosome (Ch), and complex (C) bands are indicated. Binding isotherms and K_D app_ are also shown. **B.** EMSA assays showing a two-fold dilution series (50.4-807 nM) of MeCP2 on 15-N_601_-15 nucleosomes, or chromatosomes assembled with each H1.0 tail deletion construct as indicated. DNA was methylated with linker meCpA 81 bp from the dyad. Binding isotherms and K_D app_ are also shown. **C.** Representative EMSA native gels showing a 2-fold dilution series of full-length MeCP2 on 37-N_601_-27 nucleosomes (25.2-1614 nM). Binding isotherms and K_D app_ are also shown.

Interestingly, further lengthening linkers in 37-N_601_-27 nucleosomes allowed for concurrent MeCP2 and H1.0 binding, with MeCP2 binding affinity only slightly lowered on chromatosomes versus nucleosomes (Fig. 6.C, Extended Data Fig. 9D & E)^48^. Using intermediate 16-N_603_-30 chromatosomes, with both recombinant purified H1.0 and an *ex vivo* isolated mixture of H1 isoforms, allowed for some MeCP2 binding but this was greatly reduced compared to unbound nucleosomes (Extended Data Fig. 10A-D). Overall this suggests that MeCP2 and H1 compete for proximal, but not distal, nucleosome linker DNA and require long DNA linkers in order to be co-incident.

## Discussion

MeCP2 has been the subject of intensive study, in part due to the direct genetic link to the relatively common and severe neurological disorder Rett syndrome. MeCP2 recruitment to chromatin is important for mediating its function, but has a complex binding mechanism described as both meC-dependent and independent. Here we took advantage of site-specific DNA methyltransferases and alternative DNA sequences to specifically engineer single sites of DNA methylation on a nucleosome, with the aim of clarifying disparate models of MeCP2 binding on chromatin. We identified an essential role for nucleosome linker DNA in the overall interaction of MeCP2 on nucleosomes and confirm that separable meC and DNA binding regions are important for stable interaction^35^. We observe that, facilitated by these linker DNA interactions, the MBD of MeCP2 can bind DNA methylation throughout the nucleosome (Fig. 7).

**Figure 7:**
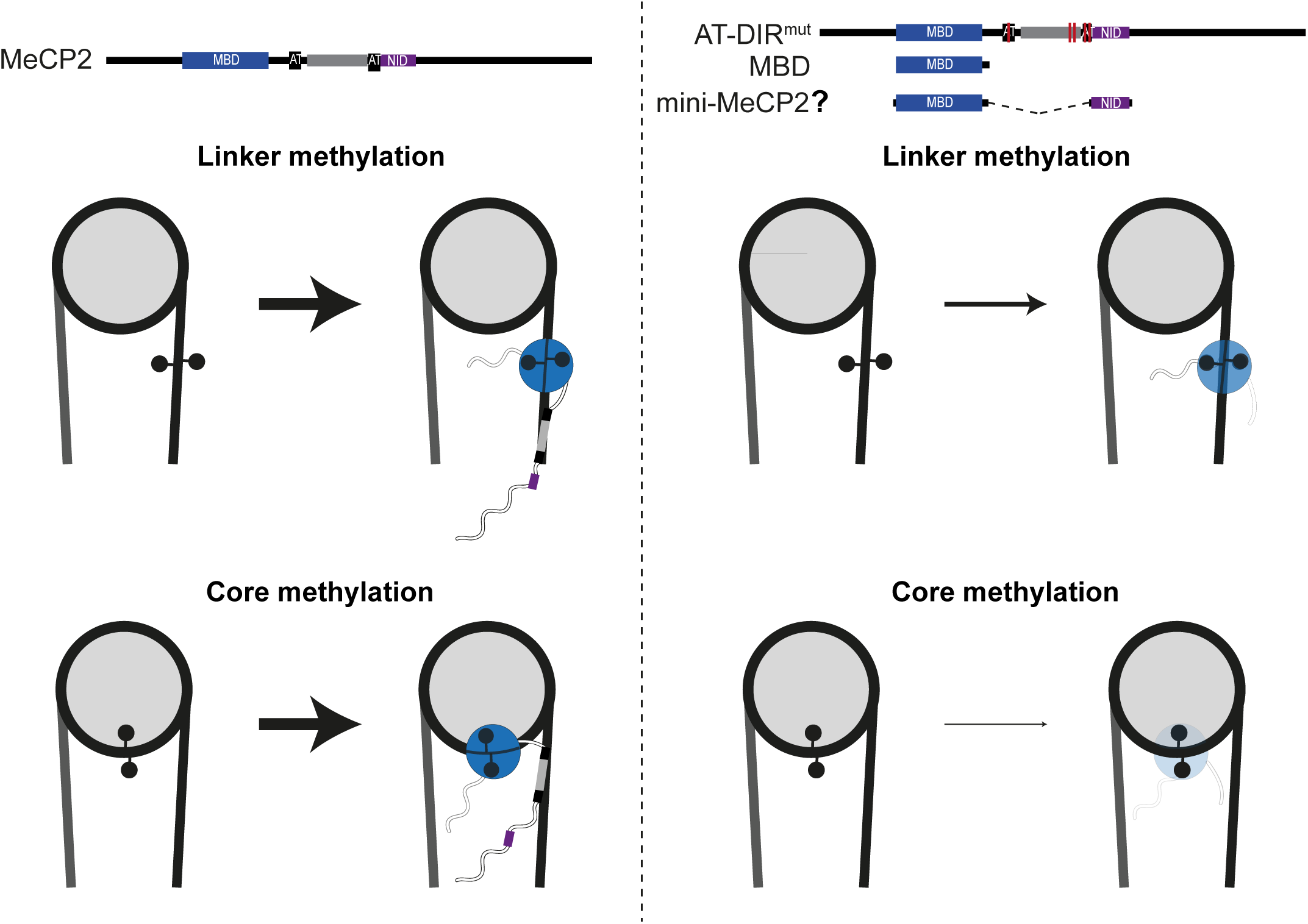
Proposed model for MeCP2 binding to nucleosomal DNA methylation. DNA methylation on linear linker DNA between nucleosome cores can be bound by variants of MeCP2 containing only the MBD, or mutations to the central region of MeCP2. However, when methylation is wrapped into nucleosomes it becomes less accessible, and the MBD alone is insufficient to bind. Additional linker DNA binding activity from central MeCP2, from two AT-hooks and a novel DIR, is required to allow nucleosomal DNA binding.

The density of meC has a large influence on MeCP2 binding^11,45,77^. Indeed, we see that the highest affinity is achieved when multiple meC are present per nucleosome (Fig. 1). However, CpG sites are under-represented in the genome, with on average 1-2 meC present per nucleosome repeat. MeCP2 levels are high in neurons, which undergo an increase in DNA methylation at non-CpG sites, roughly doubling the overall meC levels in the genome^78^. Therefore, our approach of using single meC sites within the nucleosome is representative of the meC density found *in vivo*. The relative difference in affinity for meC over unmodified nucleosomes is more notable in the case of the MBD alone, but modest in the full-length protein, due to the addition of non-specific DNA binding by the central region. We propose that *in vivo* this can manifest as both co-dependent and independent recruitment of MeCP2, depending on exact chromatin context^9,11,23,34^.

The ability of MeCP2 to bind meC at sites in the nucleosome core was surprising, although previous footprinting had suggested the possibility^43^. Some of the meC-core sites tested are in the major groove facing the histone octamer, which would be expected to occlude MeCP2 binding. Preferential binding to these DNA methylated nucleosomes therefore suggests that the nucleosome does not block access to meC to the MBD of MeCP2. Future structural data would be invaluable in explaining how the MBD can engage with meC within occluded bent core DNA.

Here we show MeCP2 binding to nucleosomal DNA methylation was dependent on nucleosome linker DNA (Fig. 2), as previously proposed^45^. Recent work suggested that MeCP2 is preferentially recruited to nucleosomes over linear DNA in single molecule assays^48^. This maybe mediated through direct histone interaction^45,48–52^, but possibly also due to the parallel nature of concurrent adjacent entry and exit DNA from a nucleosome, read in combination by the multiple DNA binding motifs found within MeCP2. At the mono-nucleosome level we find that multiple MeCP2 proteins can interact when sufficient linker DNA is available and that MeCP2 can bridge between linkers to satisfy the full DNA binding footprint (Extended data Fig. 4B & C). Whether MeCP2 preferentially binds symmetrically between both linkers or asymmetrically on a single nucleosome linker is unclear, and may be context-dependent. Furthermore how MeCP2 and H1 balance their functions is unclear. Both are present at high levels in the neuronal nucleus and have been reported to compete as well as be co-incident^11,44,48,74,79^. Intriguingly, longer linker lengths did allow the co-occurrence of MeCP2 and H1. Linker lengths are variable between cell lines and different chromatin states^80–82^, therefore distribution of nucleosomes may play a role in mediating H1 and MeCP2 engagement. We suggest that the exact positioning and abundance of meC and DNA linker-length tunes the balance between MeCP2 and H1, reflecting the changes that occur during neurodevelopment^11^.

We propose a model where DNA binding motifs in central MeCP2 interact with linker DNA, aiding the binding of the MBD to nucleosome associated DNA methylation (Fig. 7). Mutation or removal of these DNA-binding regions disrupts MeCP2 binding to nucleosomal meC. However, meC sufficiently distanced from the nucleosome does not require these additional interactions. Previous studies have suggested that regions outside of the MBD stabilise chromatin binding^57^ and the central region binds DNA^35^. We show this is required for reading of nucleosomal meC and includes previously characterised AT hooks 1 and 2, as well as a novel DNA interacting region (DIR). Primary sequence analysis mapped key residues within the DIR. While these show some overlap with the start of MeCP2’s nuclear localisation signal (NLS), the latter is not required for nuclear accumulation^83^, allowing the motif to also perform DNA-binding functions.

Separation of function mutations of conserved residues in the central region (AT-DIR^mut^) diminish MeCP2 binding and remove specificity for nucleosomal meC, without affecting the direct MBD-meC interaction. Interestingly, AT-DIR^mut^ also affects MeCP2 binding dynamics in live cells, highlighting the functional importance of the central region of MeCP2. There are few identified patient mutations in the central region compared to the clusters in the MBD and NID (Fig. 4), suggesting that this region may not be as crucial for function. However, the central region does have sections of high conservation. Perhaps, as we have seen with AT-DIR^mut^, multiple mutations of the DNA binding elements would be required to perturb function, an unlikely event in a patient setting. Nevertheless, there are patient mutations in the novel DIR region, including Rett syndrome mutant P225R. Previous work showed that an ‘mini-MeCP2’, containing only the MBD and NID domains, rescues embryonic lethality and most Rett-like phenotypes in an *Mecp2* knock-out mouse^53^. The results presented here predict that mini-MeCP2 will have reduced access the subset of DNA methylation found within the nucleosome core *in vivo*. In this context, it is interesting that the mini-MeCP2 did not fully rescue the Rett syndrome-like phenotype in mice, unlike a full-length or N-/C-terminal domain deletion construct^13,53^. We suggest that inclusion of the central region could enhance recruitment of truncated MeCP2 to chromatin, prompting a more complete rescue.

## Materials and Methods

### Generation of plasmid constructs

Histone and MeCP2 mutations were introduced either using site directed mutagenesis or direct cloning of synthesised double-strand gBlock fragments containing mutations (Integrated DNA technologies). Deletions were produced by PCR or Gibson assembly. HisMBP tagged constructs were cloned using ligation independent cloning. A summary of constructs generated in this study are listed in Supplementary Table 1.

### Histone purification

Histones were expressed in BL-21 DE3 RIL cells and purified from inclusion bodies essentially as described^84–86^.

Concentrations were determined via absorbance at 280 nm using a Nanodrop One spectrophotometer (Thermo Scientific), followed by SDS-PAGE and colloidal coomassie staining with comparison to known amounts of control proteins.

For fluorescent octamer labelling the lyophilised cysteine mutant histone H2B T115C was hydrated to 3 mg.ml^−1^ in resuspension buffer [20 mM Tris pH 7.5, 25 mM NaCl, 0.2 mM TCEP, 7 M Guanidine-HCl] for 30 mins at room temperature. Oregon-Green488 maleimide dye (AAT Bioquest) was resuspended in DMSO and added at a 1:1 molar ratio to histone H2B T115C. Samples were incubated for 2 hrs at 4°C, spiked with an equivalent volume of dye as initially added and further incubated overnight at 4°C. Labelling extent was checked by 1D intact weight ESI mass spectrometry (SIRCAMs, School of Chemistry, University of Edinburgh). Labelled histones were either used immediately for octamer assembly or flash frozen in liquid nitrogen and stored at −80°C.

H1.0 constructs were expressed and purified as previously described for the full-length protein^87^.

### Octamer assembly

Octamers were refolded as previously described^85,88^. Briefly, histones were resuspended in 20 mM Tris pH 7.5, 6 M guanidine, 10 mM DTT and mixed in a mass ratio of 1:1.4:1.6:1.6 H4, H3, H2A, H2B, and diluted to a total concentration of 2 mg.ml^−1^. The histone mixture was dialysed into 15 mM Tris pH 7.5, 2 M NaCl, 5 mM β-mercaptoethanol, 1 mM EDTA. All octamers were purified using size exclusion chromatography (HiLoad Superdex 200 16/600 or Superdex 200 Increase 10/300 GL Cytiva) in 15 mM Tris pH 7.5, 2 M NaCl, 1 mM EDTA, 5 mM β-mercaptoethanol.

H2A Lys 119 labelling with Alexa647 was performed essentially as previously described^86^ on octamers assembled with H2A K119C, H3.1 C96S C110A, H2B and H4 and desalted in an Zebaspin 7kDa column (ThermoFisher) to remove β-mercaptoethanol. 70 µM of octamer was incubated with 5 mM TCEP for 10 minutes at room temperature, 105 µM of AlexaFluor647 C2-maleimide (Invitrogen) was then added and incubated for a further 1 hour. 5 mM β-mercaptoethanol was added to quench, and the reaction desalted again as above to remove excess dye. Labelling extent was checked by measuring the 650 nm/280 nm absorbance ratio.

### MeCP2 construct purifications

Full-length untagged *H. sapien* MeCP2 (e2 isoform) constructs were expressed in Rosetta (DE3) pLysS *E. coli* cells in LB media for 3 hrs at 30°C, induced with 1 mM IPTG. Cells were pelleted at 4,000 x g for 15 mins, snap frozen in liquid nitrogen and stored at −80°C until use. Cell pellets were thawed and resuspended in lysis buffer [20 mM HEPES pH 8.0, 100 mM NaCl, 0.1% (v/v) reduced Triton X100, 100 µM PMSF, 100 µM benzamidine, 4 mM MgCl_2_, 5 µg.ml^−1^ DNase]. The suspension was nutated at 4°C for 30 mins and sonicated twice (2 secs on, 2 secs off, for a total of 20 secs at 50% amplitude). Bacterial cell debris was pelleted by centrifugation for 45 mins at 25,000 x g, 4°C. Clarified lysate was filtered through a 0.4 µm filter and buffer adjusted to 500 mM NaCl and 20 mM imidazole. A native internal stretch of Histidine residues in MeCP2 was utilised for the first step of affinity purification. Lysate was applied to a HiTrap chelating HP column (Cytiva) pre-charged with nickel ions and pre-equilibrated in Nickel A buffer [20 mM HEPES pH 8.0, 500 mM NaCl, 20 mM imidazole, 0.1% (v/v) reduced Triton-X100, 100 µM PMSF, 100 µM benzamidine]. The column was washed with 20 column volumes (CV) of Nickel A buffer and protein bulk eluted with 5 CV of Nickel B buffer [20 mM HEPES pH 8.0, 500 mM NaCl, 500 mM imidazole, 0.1% (v/v) reduced Triton-X100, 100 µM PMSF, 100 µM benzamidine]. Fractions containing the desired protein were pooled and diluted to 250 mM NaCl. Precipitate was removed by centrifugation at 4,000 x g, 4°C, for 10 mins and filtering through a 0.4 µm filter. The sample was applied to an cation exchange SP HP column (Cytiva) pre-equilibrated in SP A buffer [20 mM HEPES pH 8.0, 280 mM NaCl, 1 mM EDTA, 0.1% (v/v) reduced Triton-X100, 100 µM PMSF, 100 µM benzamidine]. The column was washed with 10 CV of SP A buffer and protein eluted across a 20 CV gradient from 280 mM to 1 M NaCl. Fractions containing the desired protein were pooled and concentrated in a 10 kDa MWCO centrifugal filter unit. The concentrated sample was applied to a HiLoad Superdex 200 16/60 column (Cytiva) pre-equilibrated in storage buffer [20 mM HEPES pH 7.5, 150 mM NaCl, 5% (v/v) glycerol, 100 µM EDTA, 5 mM β-mercaptoethanol]. Fractions containing the desired protein were pooled, concentrated as before, snap frozen in liquid nitrogen and stored at −80°C until use.

HisMBP tagged proteins were expressed and purified essentially as described above in Rosetta (DE3) pLysS as described above. Depending on purity the sample was either concentrated directly after nickel chromatography in a 10 kDa MWCO centrifugal filter unit for the final size-exclusion step or diluted to 150 mM NaCl for an additional ion-exchange step.

His-MBP-DNMT3A_1-427_ and His-MBP-DNMT3A_1-427_ K54A K56A was purified as described^72^.

To remove the His-MBP tag 1 µg of TEV protease was added to every 25 µg of protein and the sample incubated at 4°C overnight with rotation. Precipitate was removed by centrifugation at 4,000 x g, 4°C, for 10 mins and filtering through a 0.4 µm filter. The sample was then applied to an SP HP column (Cytiva) pre-equilibrated in SP A buffer [20 mM HEPES pH 7.5, 100 mM NaCl, 1 mM EDTA]. The column was washed with 5 CV of SP A buffer and protein eluted across a 20 CV gradient from 100 mM to 1 M NaCl. Fractions containing the desired protein were pooled. A 3 kDa MWCO centrifugal filter unit was used to concentrate and buffer exchange sample into storage buffer [20 mM HEPES pH 7.5, 150 mM NaCl, 5% (v/v) glycerol, 100 µM EDTA, 5 mM β-mercaptoethanol]. A sample was analysed by gel filtration on a Superdex 75 10/300 GL column (Cytiva) to check for purity. A BCA assay (Thermo) was used to determine protein concentration for DIR construct as it lacked aromatic residues. For all other MeCP2 proteins, Concentrations were determined via absorbance at 280 nm using a Nanodrop One spectrophotometer (Thermo Scientific), followed by iterative SDS-PAGE and colloidal coomassie staining with comparison to known amounts of control proteins (Supplementary Fig. 2)

### NCP reconstitution

DNA for nucleosome assembly was generated by PCR as previously described^84–86^. Sequences of DNA used are described in Supplementary Table 2, derived from Widom 601 or 603 sequences (addgene plasmid 26656 and 26658, respectively), described and used as previously^72,84,89^ or modified to alter linker length and methyltransferase recognition sites. Fluorescent dyes, biotin tags and asymmetric methyl-cytosine bases were incorporated into HPLC pure primers used in amplification PCR steps (IDT technologies). Enzymatic methylation was added to DNA at CpG sites using methyltransferase enzymes M.HhaI or M.HpaII (NEB). Reactions were set up with 1 unit of methyltransferase per µg of DNA, 640 µM S-adenosylmethionine (SAM) (NEB) and 1x rCutsmart buffer (NEB). Reactions were incubated for 3 hrs at 37°C, spiked with an equivalent volume of SAM as initially added, and further incubated overnight. Methylation state was checked by digestion with methylation sensitive HhaI or HpaII restriction enzymes (NEB). Samples were ethanol precipitated and resuspended in 10 mM Tris pH 8.0.

Nucleosomes were reconstituted essentially as described^85,88,89^. Proper assembly of wrapped nucleosomes was analysed by native PAGE and histone composition by SDS-PAGE analysis (Supplementary Fig. 1)

### DNA footprinting

DNA footprinting assays were performed essentially as described^89^. Briefly, 50 ng.µl^−1^ of 5’ 6-FAM labelled 15-N_601_-15 nucleosomes were set up in 10 µl reaction buffer (20 mM Hepes pH 7.5, 200 mM NaCl, 1 mM EDTA, 1 mM DTT). 2.5 µl each of 2 mM Ammonium Iron (II) Sulfate/4 mM EDTA, 0.1 M sodium ascorbate, and 0.12% H_2_O_2_ were simultaneously added to the sample. The reaction was stopped after 4 minutes by the addition of 100 µl STOP buffer (100 mM Tris pH 7.5, 1% glycerol, 325 mM EDTA, 0.1% SDS, 0.1 mg.ml^−1^ ProteinaseK [Thermo]) and incubated for 20 minutes at 56°C. Fragmented DNA was purified by ethanol precipitation and resuspended in 10 µl HiDi Formamide. 0.5 µl of GeneScan 500 LIZ size standard (Thermo) was also added as a size marker.

Samples were run on either a 3130xl Genetic or 3730xl DNA Analyzer, operated in accordance with the manufacturer’s instructions using the G5 dye filter set. Peaks were analysed using Thermofisher Connect Microsatellite analysis software. Peak size in base pairs were called by the Global southern method.

### Electrophoretic mobility shift assays

Nucleosomes were fluorescently labelled either on their DNA component with 5’ 6-carboxyfluorescein (5’ 6-FAM), or by the addition of Oregon-Green488 maleimide dye to H2B at residue 115 (utilising a T115C mutation). Fluorophore addition on DNA or histone was found not to not perturb MeCP2 binding. Double stranded competitor DNA was annealed using 47 bp complementary oligos incubated at equimolar ratios and heated to 95oC for 5 minutes prior to gradual cooling to room temperature.

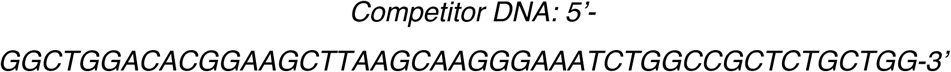

Competitor DNA was used in all nucleosome binding experiments to promote meC specificity, but absent form DNA binding experiments as highlighted in figures and figure legends.

A concentration series of purified protein (e.g. MeCP2) was incubated with 3.75 ng of fluorescent nucleosome in 12 µl of competitor EMSA buffer [20 mM HEPES pH 7.5, 75 mM NaCl, 125 mM KCl, 50 µM EDTA, 2.5 mM β-mercaptoethanol, 5% (v/v) glycerol, 333 ng.µl^−1^ BSA, 1.58 µM competitor DNA (see below)]. The reaction was incubated at room temperature for 30 mins, after which 3 µl of 5x native loading buffer [40% sucrose, 0.001% bromophenol blue] was added. A 5.2% native polyacrylamide gel was loaded with 10 µl of each sample (2.5 ng of fluorescent nucleosomes loaded) and separated at 100 V, 4°C. Gels and buffer were either made up of 0.5x TBE or 1x tris glycine. After ∼90 mins gels were imaged using a Bio-Rad ChemiDoc MP imaging system exciting using blue filtered light and recorded using filter at 532/28 emission filter. Gels were additionally stained with diamond DNA stain (Promega) and imaged using a Bio-Rad ChemiDoc MP imaging system set to 590/110 emission filter.

Unbound DNA/nucleosome bands were quantified in ImageLab (Bio-Rad) with the ‘lane and bands’ setting, and converted to “1 - relative band intensity” using:

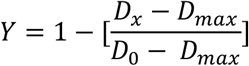

where D_x_ is the unbound band intensity at a given protein concentration X, D_0_ is the unbound band intensity at 0 µM of protein, and D_max_ is the quantification of an area equal to a band in an empty lane (equivalent to 100% bound). Data was plotted in Prism 9 (GraphPad), with a log10 x-axis, and an isotherm fitted using the method ‘specific binding with Hill slope’:

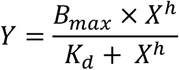

where Bmax is the maximum fraction bound (=1), Kd is the dissociation constant, h is the hill slope, and X is protein concentration. A summary of apparent Kds is reported in Supplementary Table 3. Kd apparent is reported due to the nature of competitor DNA added to the reaction and uncertainty in absolute value.

### MeCP2-H1 competition assays

The H1 concentration required to form a single clear H1 bound fluorescent nucleosome band was first identified as described for EMSAs assaying protein binding to fluorescent nucleosomes. ‘H1 mix’ is a commercial mix of bovine H1 purified from calf thymus (sigma-aldrich, 14-155) predominantly H1.4.

This determined concentration of H1 was incubated with 3.75 ng of fluorescent nucleosomes for 30 mins at 4°C in nucleosome dilution buffer [20 mM HEPES pH 7.5, 250 mM KCl, 666 ng.µl^−1^ BSA, 3.15 µM competitor DNA)], in a total volume of 6 µl. A concentration series of purified MeCP2 was then incubated with H1 bound fluorescent nucleosome in competitor EMSA buffer (20 mM HEPES pH 7.5, 75 mM NaCl, 125 mM KCl, 50 µM EDTA, 2.5 mM β-mercaptoethanol, 5% (v/v) glycerol, 333 ng.µl^−1^ BSA, 1.58 µM competitor DNA). A final reaction volume of 12 µl was incubated at room temperature for 30 mins, after which 3 µl of 5x native loading buffer was added. 10 µl of the sample (2.5 ng of fluorescent nucleosome) was loaded onto a 6% native polyacrylamide gel and separated at 100 V, 4°C. Gels and buffer were made up of 1x TG. After 120 mins the gel was imaged as before.

### MeCP2-H1-nucleosome super-shift assay

200 nM of H1.0 was incubated with 3.75 ng of fluorescent 37-N_601_27 nucleosomes for 30 mins at 4°C in nucleosome dilution buffer [20 mM HEPES pH 7.5, 250 mM KCl, 666 ng.µl^−1^ BSA, 3.15 µM competitor DNA)], in a total volume of 6 µl. 250 nM of purified MeCP2 was then incubated with H1 bound fluorescent nucleosome in competitor EMSA buffer (20 mM HEPES pH 7.5, 75 mM NaCl, 125 mM KCl, 50 µM EDTA, 2.5 mM β-mercaptoethanol, 5% (v/v) glycerol, 333 ng.µl^−1^ BSA, 1.58 µM competitor DNA), in a total volume of 12 µl for 30 mins at room temperature.

0.5 µl of Polyclonal H1.0 antibody (*ab154111*) was then added (1:25 dilution), and the 12.5 µl reaction incubated for a further 30 mins. 3 µl of 5x native loading buffer was added and 10 µl of the sample (2.4 ng of fluorescent nucleosome) was loaded onto a 5.2% native polyacrylamide gel and separated at 100 V, 4°C. Gels and buffer were made up of 1x TG. After 120 mins the gel was imaged as before.

### Bioinformatic analysis

Disease causing missense mutations from ClinVar were plotted along the length of the protein using PlotProtein^90,91^. Predicted missense severity scores were generated using Alphamissense^65^. Scores for each residue across all possible amino acids were then summed.

ClinVar mutations of interest within the MeCP2 central region were: R190H (VCV002664668.2), P217L (VCV000236302.4), P225R (VCV000143653.48), R255G (VCV000978959.2), and K266E (VCV000548706.1)

Species listed for sequence comparison are *Mus musculus* (House mouse), *Xenopus laevis* (African clawed frog), *Notamacropus eugenii* (Tammar wallaby), *Eudyptula minor* (Little penguin), *Danio rerio* (Zebrafish), and *Callorhinchus milii* (Ghost shark), with sequences retrieved from UniProt. The motif was also identified in *H. sapiens* DNMT3A and LEDGF using ScanProsite^92^ and BLASTP 2.12.0+^93^ against the uniprotkb_refprotswissprot database. The motif in LEDGF was found by literature search^70^.

### MeCP2 PFV-GAG competition

GST tagged PFV-GAG protein was purified as previously described^89^. 2 µM of MeCP2, or 500 nM of HisMBP-MeCP2_162-309_, was incubated with 3.75 ng of fluorescent nucleosomes for 30 mins at 4°C in nucleosome buffer in a total volume of 6 µl. A concentration series of GST-PFV GAG was then incubated with MeCP2 bound fluorescent nucleosomes in competitor EMSA buffer. Samples were incubated and loaded on gels as before. Gels were 5.2% native polyacrylamide.

### Microscale thermophoresis

Microscale thermophoresis (MST) measurements were performed on a Monolith NT.115 Pico instrument (NanoTemper Technologies) using standard or premium capillaries. Reconstituted H2A^K119C^ alexa647 labelled nucleosomes (at 10 nM each) were mixed 1:1 with a dilution series of MeCP2 (WT or AT-DIR^mut^) in a final buffer of [20 mM Hepes pH 7.5, 75 mM NaCl, 125 mM KCl, 0.05 mM EDTA, 5% glycerol, 0.287 mg/ml BSA, 0.02% NP-40, 2.5 mM β-mercaptoethanol, 0.3 µM competitor DNA] and incubated at room temperature for 30 min. All MST measurements were carried out at 22°C using 5% Pico-Red excitation power and ‘Medium’ MST power, with 30 s laser ON and 5 s laser OFF time. The cold phase was defined as the average signal 1 to 3 s before excitation, whilst the hot phase was defined as 1.9 to 3.9 s after excitation (Supplementary Fig. 3). Fnorm was calculated using:

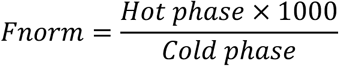

MST data was plotted in Prism 9 (GraphPad), with a log10 x-axis, and an isotherm fitted using ‘EC50 shift, X is concentration’. ΔFnorm was calculated by subtracting the bottom estimate from each Fnorm values. ΔFnorm data was fitted with an isotherm using the method ‘specific binding with Hill slope’, as performed for EMSA experiments.

### Mass photometry

Mass photometry data was collected using a TwoMP mass photometer (Refeyn), calibrated with β-amylase (56, 112, 224 kDa) and thyroglobulin (670, 1340 kDa). Movies were acquired for 2845 frames using AcquireMP software.

Nucleosome:MeCP2 complexes were mixed to a final nucleosome concentration of 50 ng.µl^−1^ in MP buffer [20 mM Na_2_HPO_4_/NaH_2_PO_4_ pH 7.5, 2 mM Hepes pH 7.5, 75 mM NaCl, 125 mM KCl, 0.5 mM EDTA, 2 mM DTT, 0.1% glycerol], and incubated on ice for 2 hours. Final molar ratios of nucleosome:MeCP2 was 1:2 for 37-N_601_-27 nucleosomes, and 1:4 for 15-N_601_-15 nucleosomes. Complexes, and nucleosomes alone, were then diluted 100-fold in dilution buffer [20 mM Na_2_HPO_4_/NaH_2_PO_4_ pH 7.5, 75 mM NaCl, 125 mM KCl, 0.5 mM EDTA, 2 mM DTT], to a final nucleosome concentration of either 3.9 nM (37-N_601_-27 nucleosomes) or 4.6 nM (15-N_601_-15 nucleosomes).

Raw data was exported from DiscoverMP software, after determining mass using calibrated standards, and plotted as a frequency distribution using Prism 9 (GraphPad) and a bin size of 5 kDa. Data was fitted with either a ‘Sum of two Gaussians’ (for 15-N_601_-15 nucleosomes) or ‘Sum of three Gaussians’ (for 37-N_601_-27 nucleosomes).

### Size Exclusion Chromatography coupled to Multi Angle Light Scattering (SEC-MALS)

Size-exclusion chromatography (AKTA PURE25^TM^; Cytiva) coupled with UV, static light scattering and refractive index (RI) detection (Viscotec SEC-MALS 20 and Viscotek RI Detector VE3580; Malvern Instruments) were used to determine the absolute molecular mass of untagged MeCP2 DIR (MeCP2_205-257_) in solution. A 100 µL injection of 2.11 mg.mL^−1^ (375 µM) untagged MeCP2_205-257_ was run on a calibrated Superdex-75 Increase 10/300 GL (Cytiva) size exclusion column pre-equilibrated in 20 mM HEPES, pH 7.5, 150 mM NaCl, 0.1 mM EDTA, 2.5% (v/v) glycerol at 22 °C with a flow rate of 1.0 mL.min^−1^. Light scattering, RI and A280 (protein elution from the chromatography system was monitored at 230 nm due to the protein construct having no aromatics) were analysed by a homo-polymer model (OMNISEC software, v5.1; Malvern Instruments) using the following parameters for MeCP2 DIR (MeCP2_205-257_): dA/dc = 0.01 AU.mL^−1^.mg^−1^, dn/dc = 0.185 mL.g^−1^ and buffer RI value of 1.3362. Peak fractions were run on a 17% SDS-PAGE gel for analysis.

### Surface Plasmon Resonance (SPR)

Nucleosomes wrapped with 37-N_601_-27 biotinylated DNA were immobilised on streptavidin sensor surfaces at 20 nM, at 25°C at a flow rate of 5 µL.min^−1^ on a BIAcore T200 instrument (Cytiva) equilibrated in *SPR Buffer* [20 mM HEPES, pH 7.5; 200 mM KCl; 1 mM EDTA; 0.5 mM DTT; 0.02% NP-40; 0.33 mg.mL^−1^ BSA], Flow cell 1 (blocked with 20 nM D-biotin in *SPR Buffer*) was used as the bulk reference surface, while 3x meCpG, −1 meCpG or unmethylated nucleosomes were separately immobilised on flow cells 2,3 and 4, respectively, to between 1,540 and 1,580 RU. A 2-fold dilution series (12.5 - 200 nM) of WT or AT-DIR^mut^ MeCP2 protein, supplemented with 50 ng.µl^−1^ of competitor DNA, was injected at 30 µL.min^−1^, in a single-cycle kinetic experiment each with a 60 second contact time, a 60 second dissociation phase, and a final 600 second dissociation. A 30 second injection, at 30 µL.min^−1^ of 20 mM HEPES (pH 7.4); 1.5 M KCl; 1 mM EDTA, 0.5 mM DTT, 0.02% P20; 0.33 mg/mL BSA was used to regenerate the surface between cycles.

### Crosslinking mass spectrometry

Complexes of MeCP2 bound to methylated 175 bp nucleosomes (2.5:1 molar ratio) were crosslinked using a concentration range of photo-reactive sulfo-SDA (sulfosuccinimidyl 4,4’-azipentanoate) (Thermo Fisher) in a w/w ratio of 1:0.25-1.5 (nucleosome: sulfo-SDA) in crosslink buffer [20 mM HEPES pH7.5, 250 mM NaCl, 1 mM EDTA, 1 mM DTT]. A total reaction volume of 30 µl, containing 7.5 µg of nucleosome, was incubated for 2 hrs on ice prior to UV irradiation at 365nm in a CL-1000L UV cross-linker (Spectrum) for 20 mins. The reaction was immediately quenched with 40 mM ammonium bicarbonate. The crosslinked sample was separated on an 3-8% gradient nuPAGE gel (Invitrogen) and bands running at a higher molecular weight than MeCP2 were excised. Protein gel bands were reduced with 10 mM TCEP for 30min at 37°C, alkylated with 55 mM iodoacetamide for 20min at room temperature and digested using 13 ng.μl^−1^ trypsin (Promega) overnight at 37°C. Digested peptides were desalted using C18-StageTips^94^ for LC-MS/MS analysis.

LC-MS/MS analysis was performed using Orbitrap Fusion Lumos (Thermo Fisher Scientific) with a “high/high” acquisition strategy. The peptide separation was carried out on an EASY-Spray column (50 cm × 75 μm i.d., PepMap C18, 2 μm particles, 100 Å pore size, Thermo Fisher Scientific). Mobile phase A consisted of water and 0.1% v/v formic acid. Mobile phase B consisted of 80% v/v acetonitrile and 0.1% v/v formic acid. Peptides were loaded at a flow rate of 0.3 μl/min and eluted at 0.2 μl/min using a linear gradient going from 2% mobile phase B to 40% mobile phase B over 139 min (each sample has been running three time with different gradient), followed by a linear increase from 40% to 95% mobile phase B in 11 min (160 min total run time). The eluted peptides were directly introduced into the mass spectrometer. MS data were acquired in the data-dependent mode with 3 seconds acquisition cycle. Precursor spectrum was recorded in the Orbitrap with a resolution of 120,000. The ions with a precursor charge state between 3+ and 8+ were isolated with a window size of 1.6 m/z and fragmented using high-energy collision dissociation (HCD) with collision energy 30. The fragmentation spectra were recorded in the Orbitrap with a resolution of 30,000. Dynamic exclusion was enabled with single repeat count and 60 seconds exclusion duration.

Peak lists were generated with ProteoWizard (version 3.0.24283)^95^, and cross-linked peptides were matched to spectra using Xi software (version 1.8.4.1)^96^; Xi search) with in-search assignment of monoisotopic peaks (Lenz et al, 2018). MS1 accuracy was set to 3 ppm and MS2 accuracy to 10 ppm. Trypsin was the protease of choice allowing four missed cleavages and SDA was chosen from the crosslinker list. Carbamidomethylation of cysteine was chosen as a fixed modification and oxidation of methionine was chosen as variable modification. TR False discovery rate was computed using XiFDR and results reported at 1% residue level false discovery rate^97^. Our in-house protein database was used for the searches, containing MeCP2, and histones H2A, H2B, H3.1 and H4 (all *Homo sapiens*).

### Cell culture and transfection

NIH-3T3 mouse fibroblasts (ECACC, 93061524) were cultured in Dulbecco’s Modified Eagle Medium (DMEM; Gibco ref. 41966029) supplemented with 10% fetal bovine serum and were grown at 37°C with 5% carbon dioxide. For imaging, 1.5X10^5^ cells were plated and cultured directly on polymer coverslips (iBidi cat. 81156) with gelatine coating and the appropriate culture conditions. NIH-3T3 cells were transfected with 2µg of wild-type eGFP-MeCP2 or mutant versions (Δ163-271, AT-DIR^mut^, R133C) using Lipofectamine 2000 (Thermo Fisher Scientific cat. 11668019), following the manufacturer’s protocol.

### Live cell imaging

Live cells were imaged 24 hours after transfection. NucBlue Live Cell (ThermoFisher cat. R37605) stain was added prior to imaging, as directed by the manufacturer, using a Zeiss LSM 880 confocal microscope with an Airyscan module and environmental chamber at 37°C with 5% carbon dioxide. Images were acquired using Zeiss ZEN software (black edition) and processed using Fiji software.

### Fluorescence recovery after photo-bleaching (FRAP)

Live cells were analysed the day after transfection and FRAP was performed using a Zeiss LSM 880 confocal microscope, equipped with Airyscan module and environmental chamber at 37°C with 5% carbon dioxide. For each cell, eGFP-MeCP2 signal was imaged every 1sec for 400sec with five images recorded before bleaching at a selected MeCP2-enriched spot (FRAP spot) with 100% laser power.

FRAP analysis of three independent transfection experiments was performed using a custom macro with Fiji software (https://doi.org/10.5281/zenodo.2654601). Mean pre-bleach fluorescence was estimated per cell to control for transfection efficiency (Extended Data Fig. 8A). Fluorescence was measured at the bleached MeCP2 spot (FRAP spot), as well as a non-bleached MeCP2 spot (control spot) to account for photobleaching during the experiment. Additionally, the fluorescence outside of transfected cells was measured as background. The first time point (T0) was defined as the first post-bleach image. For each time point, the FRAP fluorescence signal was normalised to fluorescence values before photobleaching, as described in the equation below:

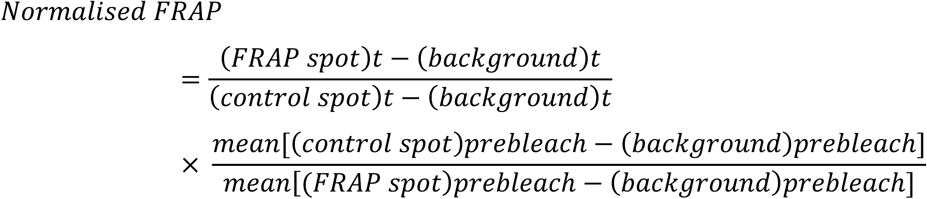

A “Two phase association” model (nonlinear regression) was used to fit experimental data using the software GraphPad Prism 10. The plateau (fluorescence at the last time point of the FRAP experiment), corresponding to the mobile fraction, was interpolated from the fitted curve. The T1/2 (time to recover 50% of fluorescence at plateau) was interpolated from the fitted curve. Data was plotted using GraphPad Prism 10.

## Data Availability

Crosslinking mass spectrometry data has been uploaded to PRIDE database under identifier PXD064826.

## Acknowledgements

We thank Nick Gilbert, Owen Davies and members of the Wilson lab for helpful discussions. MDW’s work is supported by the Wellcome Trust [210493], Medical Research Council (T029471/1). JAW’s work was supported by integrative cellular mechanisms PhD studentship (218470). A.B.’s research is funded by a Wellcome Investigator Award (#107930) and grants from the Simons Initiative for the Developing Brain and the Rett Syndrome Research Trust. This work was supported by the Edinburgh Protein Production Facility (EPPF), which received funding from a core grant (203149) to the Wellcome Centre for Cell Biology at the University of Edinburgh. We thank Gauri Déak for the PFV-GAG protein. We are grateful to the Rappsilber lab for access to the Xi server for mapping protein crosslinks. We would like to thank Prof. Louise E. Horsfall’s group at University of Edinburgh for access to MST equipment, which was funded by the EPSRC (EP/N026519/1), and Michael Capeness for help and advice on these experiments. This work was supported by funding for the Wellcome Discovery Research Platform for Hidden Cell Biology [226791] and we gratefully acknowledge support from the Light Microscopy, Proteomics and Structural Biology cores. The proteomics facility is supported by Multi-User Equipment grant (108504) and we are grateful to Christos Spanos for advice with experiment design and PRIDE submission. We thank Logan Mackay in SIRCAMS school of chemistry, university of Edinburgh for 1D Mass Spec analysis. We also thank the Technical Services Sequencing Facility at The Institute of Genetics and Cancer, University of Edinburgh for processing hydroxyl radical footprinting samples. We are grateful to Pamela Brown at the Institute for Regeneration and Repair Biomolecular and Assay Core for access and help with Mass photometry analysis.

## Competing Interests

The authors declare no competing interests.

## Extended Data Figure Legends

**Extended Data Figure 1: meCpG position on Widom601 nucleosomes.**

**A.** Location of meCpG dinucleotides −1 bp (grey) and −61 bp (brown) from the dyad, shown both as cartoons and on the crystal structure of a Widom 601 nucleosome (PDB: 3LZ0^98^). SHL locations are marked.

**B.** Hydroxyl radical footprinting of 15-N_601_-15 nucleosomes either unmethylated (blue), or methylated with meCpG at position −1 (grey) or −61 (brown). A 10 bp periodicity of protected bases is indicative of histone-DNA interactions. Location of methylated DNA on the footprinting traces is indicated below.

**Extended Data Figure 2: MeCP2 can bind DNA methylation along the nucleosome linker.**

**A.** Representative EMSA native gels (3 repeats) showing a 2-fold dilution series of MBD (left) (26.8-428 nM) and a 1.5-fold dilution series of MeCP2 (right) (56.7-287 nM) on 16-N_603_-30 nucleosomes. Nucleosomes were either methylated with meCpA at position 91 (red), position −81 (orange), or unmethylated (blue). Binding isotherms and K_D app_ are also shown.

**B.** Representative EMSA native gels (3 repeats) showing a 1.5-fold dilution series of MeCP2 on 16-N_603_-30 nucleosomes (56.7-287 nM) containing meCpG at positions 91 (red) and 81 (orange), or unmethylated (blue). Binding isotherms and K_D app_ are also shown.

**C.** Mass photometry analysis for the complexes formed by MeCP2 on 37-N_601_-27 nucleosomes (left), or on 15-N_601_-15 nucleosomes (right). MP of both nucleosomes alone is also shown for comparison. Based on expected mass, peak (a) corresponds to unbound nucleosome, whilst (b)-(c) are 1:1 and 1:2 nucleosome:MeCP2 complexes. In all examples peaks were also observed measuring at around 100 kDa (d), which likely corresponds to unwrapped free DNA.

**Extended Data Figure 3: MeCP2 binds linker DNA alongside the H3 N-terminal tail.**

**A.** Circular representation of Sulfo-SDA crosslinks between MeCP2 and histones on nucleosome containing 91 meCpA (left), −1 meCpG (middle), and −61 meCpG (right). Graphics of 15-N_601_-15 nucleosomes are shown below, with crosslinked histone residues highlighted by darker shading. Each graphic is split into crosslinks to the central region of MeCP2 (left) and crosslinks to the MBD (right).

**B.** Representative EMSA native gels (3 repeats) showing a two-fold dilution series (13.4-1713 nM) of MBD on 15-N_601_-15 nucleosomes. Nucleosomes were either wild-type (black), H3 tailless (light blue), or H3K27me3 (dark blue). Binding isotherms and K_D app_ are also shown.

**C.** Representative EMSA native gels (3 repeats) showing a 1.5-fold dilution series (25.2-191 nM) of MeCP2 on 0-N_603_-30 nucleosomes. Nucleosomes were either wild-type with 91 meCpA (red), H3 tailless with 91 meCpA (light red), wild-type unmethylated (dark blue), or H3 tailless unmethylated (light blue). Binding isotherms and K_D app_ are also shown.

**Extended Data Figure 4: Nucleosome linker length specifies MeCP2 binding to nucleosomal DNA methylation.**

**A.** EMSA native gels showing a two-fold dilution series (31.3-1000 nM) of MeCP2 binding to 16-N_603_-30 (dark blue), 15-N_601_-15 (light blue) and NCPs N_601_ (grey) nucleosomes and their corresponding DNA. Binding to DNA is shown on the left of each gel, nucleosomes on the right.

**B.** Representative EMSA native gels (3 repeats) showing a two-fold dilution series (50.4-807 nM) of MeCP2 on 0-N_601_-15 nucleosomes. Nucleosomes were methylated with meCpA at position −81 (pink), or unmethylated (light blue). Binding isotherms and K_D app_ are also shown. Data was plotted alongside data for binding to 15-N_601_-15 nucleosomes (dark red, dark blue) (Fig. 2) repeated here for comparison.

**C.** Representative EMSA native gels showing a two-fold dilution series (25.2-404 nM) of MeCP2 on 0-N_603_-30 and 16-N_603_-30 nucleosomes (3 repeats). Nucleosomes were methylated with meCpA at −91 bp (light/dark red), or unmethylated (light/dark blue). Binding isotherms and K_D app_ are also shown.

**D.** A model for linker DNA dependent binding events of MeCP2 on nucleosomes. MeCP2 binds once to 15-N_601_-15 nucleosomes. Both linkers are needed for optimal binding. While the MBD likely binds the meCpA site on one linker, a region of MeCP2 binding to the other linker is hypothesised. There are two binding events on 16-N_603_-30 nucleosomes. MeCP2 first binds to the 30 bp linker containing a meCpA site, likely through an interaction with the MBD. The addition of a 16 bp second linker allows the DNA-mediated binding of a second MeCP2 at high protein concentrations.

**Extended Data Figure 5: MeCP2 weakly contacts the nucleosome acidic patch.**

**A.** Representative EMSA native gels (2 repeats) showing a dilution series (375, 500, 750, 1000, 1500 nM) of MeCP2_162-309_ on N_601_ nucleosomes. Nucleosomes were either wild-type (WT) (red) or acidic patch mutant at key negatively charged residues in H2A and H2B (blue). Binding isotherms and K_D app_ are also shown.

**B.** Representative EMSA native gels (3 repeats) showing a two-fold dilution series (125-2000 nM) of MeCP2_162-309_ on 15-N_601_-15 nucleosomes. Nucleosomes were either wild-type (WT) (red) or acidic patch mutant (blue). Binding isotherms and K_D app_ are also shown.

**C.** EMSA native gel showing a two-fold dilution series (7.8-2000 nM) of acidic-patch binding PFV-GAG protein on 16-N_603_-30 nucleosomes either alone (top), pre-bound with 2 µM of MeCP2 (middle), or pre-bound with 500 nM of HisMBP-MeCP2_162-309_ (bottom). The lane with equimolar amounts of MeCP2 and PFV-GAG is marked (*), suggesting MeCp2 and PFV Gag can co-exist on nucleosomes.

**D.** Representative EMSA native gels (3 repeats) for the additional data plotted in Fig. 3B. showing a two-fold dilution series of MeCP2 R133G (27.2-435 nM) and MeCP2_162-309_ (31.3-500 nM) construct on 16-N_601_-30 nucleosomes. Nucleosomes were either methylated with meCpA at position −91 (red) or unmethylated (blue). A control HisMBP EMSA was also performed (2 repeats) (31.3-500 nM).

**E.** Representative EMSA native gels (3 repeats) for the additional data plotted in Fig. 3B. showing a two-fold dilution series of MeCP2 R133G (27.2-435 nM), MeCP2_162-309_ (31.3-500 nM) and MBD (26.8-428 nM) construct on 15-N_601_-15 (light blue) and N_601_ (grey) unmethylated nucleosomes.

**Extended Data Figure 6: Further biochemical analysis of the DIR of MeCP2.**

**A.** Representative EMSA native gels (2 repeats) showing a two-fold dilution series (125-16000 nM) of His-MBP tag alone control incubated with 40 bp (dark green) and 20 bp (light green) dsDNA.

**B.** Representative EMSA native gels (3 repeats) showing a two-fold dilution series (125-16000 nM) of MeCP2_205-257_ on 40 bp (dark green) and 20 bp (light green) DNA. Binding isotherms and K_D app_ are also shown. No competitor DNA was used in this assay.

**C.** Representative EMSA native gels (2 repeats) assaying MeCP2_205-257_ (63, 250, 1000 nM) binding to a panel of varied DNA sequences with annotated GC content (red-yellow). Quantification of the free DNA bands at each concentration, for every DNA target, is plotted. No competitor DNA was used in this assay.

**D.** (left) Size-exclusion chromatography trace of purified tag cleaved MeCP2_205-257_. 280 nm (green) and 230 nm (red) traces are shown. Note that this construct does not absorb at 280 nm. Peak fractions were run on a 17 % SDS-PAGE gel and stained for protein (middle). Tag cleaved MeCP2_205-257_ is 5.6 kDa. SEC-MALS analysis of the peak was also performed (right). Refractive index (mV) and calculated molecular weight (Da) are shown. Molecular weight (Mw) determined by amino acid composition (calculated), size-exclusion chromatography elution volume (elution apparent) and SEC-MALS measurement (experimental) are also shown below.

**E.** Representative EMSA native gels (3 repeats) showing a two-fold dilution series (15.6-4000 nM) of His-MBP-DNMT3A_1-427_ variants binding to 40 bp dsDNA. Binding isotherms and K_D app_ are also shown. No competitor DNA was used in this assay.

**Extended Data Figure 7: Disruption of DNA interactors in central MeCP2 disturbs binding to nucleosomes.**

**A.** SPR trace of MeCP2 WT (solid) and AT-DIR^mut^ (dashed) binding to 37-W_601_-27 nucleosomes. Nucleosomes were methylated with meCpG at three positions (red), a single meCpG at position −1 bp (grey), or unmethylated (blue). A 2-fold concentration series of protein (12.5-200 nM) was injected sequentially. Comparisons of 25 nM protein association/dissociation phases (160-300 seconds) are enlarged(right).

**B.** Representative EMSA native gels (3 repeats) showing a 1.5-fold dilution series of MeCP2 WT (56.7-656 nM) and AT-DIR^mut^ (R188G, R190G, K254N, K256N, R268Q) (56.3-641 nM) on 15-N_601_-15 nucleosomes. Nucleosomes were methylated with meCpG either at position - 61 (brown) position −1 (grey), or unmethylated (blue). Binding isotherms and K_D app_ are also shown.

**C.** Representative EMSA native gels (3 repeats) showing a 1.5-fold dilution series of MeCP2 WT (56.7-287 nM) and AT-DIR^mut^ (R188G, R190G, K254N, K256N, R268Q) (56.3-285 nM) on 16-N_603_-30 nucleosomes. Nucleosomes were methylated with meCpA either at position 91 (red) position 81 (orange), or unmethylated (blue). Binding isotherms and K_D app_ are also shown.

**Extended Data Figure 8: MeCP2 constructs localise to pericentric heterochromatin foci in mouse fibroblasts.**

(Left) Live cell imaging of NIH3T3 mouse fibroblast cells transfected with each eGFP labelled MeCP2 construct as indicated. Hoechst staining was used to visualise DNA. 5 µm scale bars are shown. (Right) Scatter plot of the mean fluorescence of each cell before bleaching and FRAP analysis. Median fluorescence of each construct is shown.

**Extended Data Figure 9: Formation of H1.0 chromatosomes.**

**A.** SDS-PAGE loaded with 1 µg of each H1.0 construct.

**B.** EMSA native gel showing a two-fold dilution series (15.6-500 nM) of purified H1.0 protein on 15-N_601_-15 nucleosomes with either −81 meCpA (red), −1 meCpG (grey), −61 meCpG (brown), or unmethylated (blue).

**C.** EMSA native gel showing a two-fold dilution series (31.3-1000 nM) of H1.0 deletion proteins on 15-N_601_-15 nucleosomes with meCpA at position −81.

**D.** EMSA native gel showing a two-fold dilution series (3.9-2000 nM) of H1.0 proteins on 37-W_601_-27 nucleosomes with either three meCpG sites (red) or unmethylated (blue).

**E.** Super-shift EMSA native gel to probe whether H1 and MeCP2 can be coincident on nucleosomes identifying nature of observed bands. 37-W_601_-27 nucleosomes bound by H1.0 (100 nM) and MeCP2 (250 nM) were incubated. Polyclonal H1.0 antibody (*ab154111*) was added in a 1:25 dilution to induce super-shifting.

**Extended Data Figure 10: H1 variants also compete with MeCP2 for nucleosome binding.**

**A.** EMSA native gel showing a two-fold dilution series (3.9-2000 nM) of purified H1.0 protein (left) or commercially bought H1 mix protein (sigma-aldrich, 14-155) (right) on 16-N_601_-30 nucleosomes with either −91 meCpA (red) or unmethylated (blue).

**B.** EMSA native gels (2 repeats) showing a two-fold dilution series (12.6-404 nM) of MeCP2 on 16-N_601_-30 nucleosomes (left) or chromatosomes (right). DNA was either methylated with −91 meCpA (red), or unmethylated (blue). The concentration point at which MeCP2 and H1 were added in equimolar amounts is marked (*).

**C.** Repetition of (b) with commercial H1 mix.

**D.** EMSA native gels showing a two-fold dilution series (31.3-250 nM) of H1.0 on 16-N_603_-30 nucleosomes either alone (left) or pre-bound with 250 nM of MeCP2 (right).

## Supplementary Figure legends

**Supplementary Figure 1: Quality control of *in vitro* reconstituted nucleosomes and recombinant MeCP2.**

**A.** SDS-PAGE gels loaded with 1 µg of each purified construct. MBD (left) and full-length MeCP2(right). The 52 kDa MeCP2 protein runs at around 72 kDa.

**B.** Size exclusion chromatography trace of assembled OG488 labelled H2B^T115C^ octamers produced three distinct peaks. OG488 dye was followed by absorbance at 498 nm (yellow, dashed), protein at 280 nm (green, solid) and DNA at 260 nm (blue, solid). Peak fractions were run on a 17% SDS-PAGE gel, imaged for fluorescence, and stained for protein. Peak 1 primarily contained excess H2B, peak 2 the histone octamer, and peak 3 excess H2A H2B dimer.

**C.** Restriction digest reaction of methylated and unmethylated 37-N_601_-27 and 15-N_601_-15 DNA by M.HhaI or M.HpaII. A lack of digestion upon addition of HhaI or HpaII restriction enzyme is indicative of complete methylation. Final expected DNA sizes upon complete digestion are marked.

**D.** (Left) Representative native polyacrylamide gel of H2B^T115C^ OG488 labelled octamers wrapped with 37-N_601_-27 DNA. The gel was imaged for fluorescence before being stained for DNA. A band shift is indicative of nucleosome formation (N). Unwrapped DNA is included as a control (D). (Right) Representative native polyacrylamide gel of H2B^T115C^ OG488 labelled octamers wrapped with 15-N_601_-15 DNA, and unlabelled octamers wrapped with FAM labelled 15-N_601_-15 DNA. The gel was imaged for fluorescence before being stained for DNA. A band shift is indicative of nucleosome formation (N). Nucleosomes were purified from excess DNA (P) by precipitation with PEG6000. Unwrapped DNA (D) is included as a control.

**E.** Representative EMSA native gel (2 repeats) showing the effect of a two-fold dilution series of competitor DNA (0.4, 0.8, 1.6, 3.2 µM) on the binding of full-length MeCP2 (250 nM) to 16-N_603_-30 nucleosomes. Nucleosomes were either methylated with meCpG at position 1 (red), or unmethylated (blue).

**Supplementary Figure 2: Coomassie SDS-PAGE gels of purified proteins**

**A.** 17% SDS-PAGE of modified H3 tailless nucleosomes used in Extended Data Fig. 3B.

**B.** 17% SDS-PAGE of modified H3 tailless and H3K27me3 nucleosomes used in Extended Data Fig. 3C.

**C.** 12% SDS-PAGE gel loaded with 1 µg of each purified HisMBP tagged construct (Fig. 4).

**D.** 12% SDS-PAGE gel loaded with 1 µg of purified wild-type and K254N/K256N HisMBP-MeCP2_205-257_ construct (Fig. 4).

**E.** 12% SDS-PAGE gel loaded with 0.5 and 0.25 µg of wild-type, R133G and AT-DIR^mut^ (R188G, R190G, K254N, K256N, R268Q) full-length MeCP2.

**Supplementary Figure 3: MST traces**

Raw MST traces of MeCP2 WT and AT-DIR^mut^ titrated against H2AK119C-alexa647 labelled 15-N_601_-15 nucleosomes. Cold and hot regions are highlighted in blue and red. Initial, thermophoresis, and recovery phases are indicated by dotted lines.

**Supplementary Table 1:** Expression constructs used in this study.

| Constructs for MeCP2 *E.coli* expression | Source |
| --- | --- |
| MeCP2 FL | 99 |
| MeCP2 R133G | This study |
| MeCP2 K254N,K256N,R188G,R190G,R268Q | This study |
| MeCP2 77-167 6xHis | 100 |
| 6xHisMBP MeCP2 162-309 | This study |
| 6xHisMBP MeCP2 205-257 | This study |
| 6xHisMBP MeCP2 205-257 K254N K256N | This study |
| 6xHisMBP MeCP2 205-309 | This study |
| 6xHisMBP MeCP2 258-309 | This study |
| 6xHisMBP MeCP2 272-309 | This study |
| Constructs for MeCP2 NIH3T3 expression | Source |
| MeCP2 FL | 60 |
| MeCP2 R133C | 101 |
| MeCP2 Δ163-271 | This study |
| MeCP2 K254N,K256N,R188G,R190G,R268Q | This study |
| Constructs for histone *E.coli* expression | Source |
| H1.0 TEV-6xHis | This study |
| H1.0 1-97 TEV-6xHis | This study |
| H1.0 25-194 TEV-6xHis | This study |
| H1.0 25-194 TEV-6xHis | This study |
| unmodified H3.1 (No cys) | 86 |
| H3.1 K27C | 72 |
| H2A | Addgene |
| H2A E61A E91A E92A | 102 |
| H2A K119C | 84 |
| H2B | Addgene |
| H2B T155C | This study |
| H2B E113A | 102 |
| H4 | Addgene |
| H3 A25C 25-135 | 102 |
| Constructs for additional protein *E.coli* expression | Source |
| 6xHisGST PFV GAG 534-557 | 89 |
| 6xHisMBP DNMT3A 1-427 | 72 |
| 6xHisMBP DNMT3A 1-427 K54A K56A | 71 |
| Constructs for nucleosomal DNA | Source |
| N<sub>601</sub> | 89 |
| 15-N<sub>601</sub>-15 | 84 |
| N<sub>601</sub>-15 | This study |
| 37-N<sub>601</sub>-27 | 72 |
| 16-N<sub>603</sub>-30 | This study |
| 16-N<sub>603</sub>-30 | This study |
| 16-N<sub>603</sub>-30 +81 CG | This study |
| 16-N<sub>603</sub>-30 +91 CG | This study |

**Supplementary Table 2:**
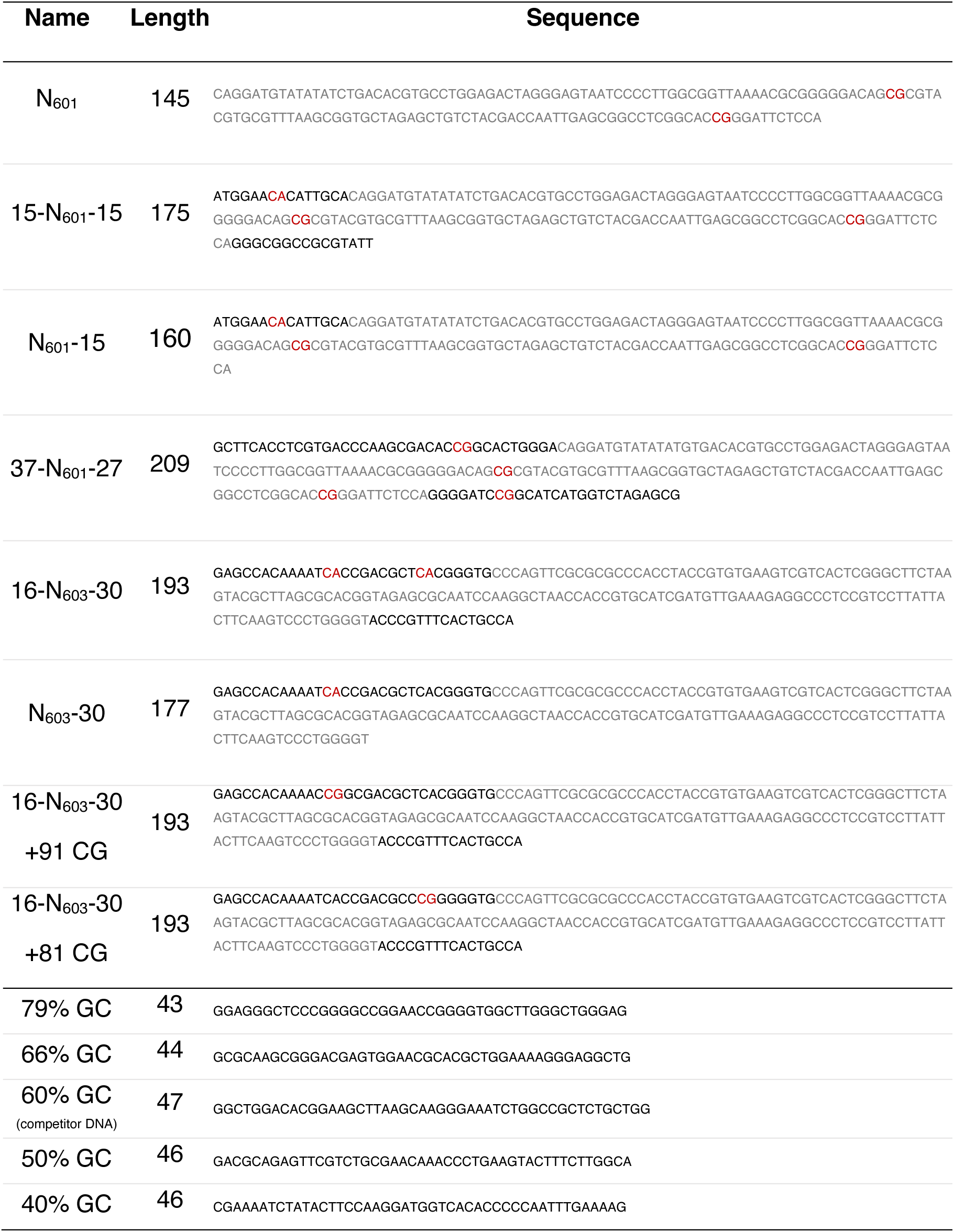
DNA sequences used for wrapping nucleosomes and binding assays.

**Supplementary Table 3:** Summary of affinity measurements assessed by EMSA assays.

| Protein construct | DNA length | Substrate type | Methylation state | K<sub>D</sub><br>app |  |  | h |  |  |
| --- | --- | --- | --- | --- | --- | --- | --- | --- | --- |
| FL MeCP2 | 37-N601-27 | Nucleosome | 3x meCpG | 37.5 | ± | 1.5 | 1.7 | ± | 0.1 |
| FL MeCP2 | 37-N601-27 | Nucleosome | position -1 meCpG | 72.1 | ± | 3.2 | 1.8 | ± | 0.1 |
| FL MeCP2 | 37-N601-27 | Nucleosome | unmethylated | 137.4 | ± | 6.4 | 2.1 | ± | 0.2 |
| FL MeCP2 | 15-N601-15 | Nucleosome | position -1 meCpG | 130.1 | ± | 3.3 | 2.6 | ± | 0.1 |
| FL MeCP2 | 15-N601-15 | Nucleosome | position -61 meCpG | 137.4 | ± | 3.8 | 3.7 | ± | 0.3 |
| FL MeCP2 | 15-N601-15 | Nucleosome | unmethylated | 220.5 | ± | 7.5 | 3.7 | ± | 0.4 |
| FL MeCP2 | N601 | Nucleosome | position -1 meCpG | N.D. |  |  | N.D. |  |  |
| FL MeCP2 | N601 | Nucleosome | position -61 meCpG | N.D. |  |  | N.D. |  |  |
| FL MeCP2 | N601 | Nucleosome | unmethylated | N.D. |  |  | N.D. |  |  |
| FL MeCP2 | 16-N603-30 | Nucleosome | position 91 meCpA | 57.1 | ± | 1.4 | 2.8 | ± | 0.2 |
| FL MeCP2 | 16-N603-30 | Nucleosome | position 81 meCpA | 65.3 | ± | 3.3 | 3.1 | ± | 0.4 |
| FL MeCP2 | 16-N603-30 | Nucleosome | unmethylated | 94.9 | ± | 2.7 | 3.6 | ± | 0.3 |
| FL MeCP2 | 16-N603-30 | Nucleosome | position 91 meCpG | 71.3 | ± | 1.4 | 3.5 | ± | 0.2 |
| FL MeCP2 | 16-N603-30 | Nucleosome | position 81 meCpG | 65.5 | ± | 1.6 | 2.9 | ± | 0.2 |
| FL MeCP2 | 16-N603-30 | Nucleosome | unmethylated | 137.7 | ± | 4.0 | 3.4 | ± | 0.3 |
| FL MeCP2 | 15-N601-15 | Nucleosome | position 81 meCpA | 140.8 | ± | 11.3 | 2.4 | ± | 0.4 |
| FL MeCP2 | 0-N601-15 | Nucleosome | position 81 meCpA | 357.1 | ± | 19.7 | 1.5 | ± | 0.1 |
| FL MeCP2 | 0-N601-15 | Nucleosome | unmethylated | 419.0 | ± | 42.8 | 1.5 | ± | 0.2 |
| FL MeCP2 | 0-N603-30 | Nucleosome | position 91 meCpA | 61.4 | ± | 2.2 | 2.6 | ± | 0.2 |
| FL MeCP2 | 0-N603-30 | Nucleosome | unmethylated | 101.7 | ± | 6.3 | 3.4 | ± | 0.8 |
| FL MeCP2 | 0-N603-30 | Nucleosome,<br>H3tailless | position 91 meCpA | 47.0 | ± | 1.1 | 1.8 | ± | 0.1 |
| FL MeCP2 | 0-N603-30 | Nucleosome,<br>H3tailless | unmethylated | 86.9 | ± | 2.3 | 3.4 | ± | 0.3 |
| FL MeCP2 | 15-N601-15 | H1.0<br>Chromatosome | position -1 meCpG | N.D. |  |  | N.D. |  |  |
| FL MeCP2 | 15-N601-15 | H1.0<br>Chromatosome | position -61 meCpG | N.D. |  |  | N.D. |  |  |
| FL MeCP2 | 15-N601-15 | H1.0<br>Chromatosome | unmethylated | N.D. |  |  | N.D. |  |  |
| FL MeCP2 | 37-N601-27 | H1.0<br>Chromatosome | 3x meCpG meCpG | 46.0 | ± | 2.4 | 1.4 | ± | 0.1 |
| FL MeCP2 | 37-N601-27 | H1.0<br>Chromatosome | position -1 meCpG | 80.4 | ± | 7.7 | 1.4 | ± | 0.2 |
| FL MeCP2 | 37-N601-27 | H1.0<br>Chromatosome | unmethylated | 164.5 | ± | 16.8 | 2.2 | ± | 0.4 |
| FL MeCP2 | 15-N601-15 | H1.0<sub>25-194</sub><br>Chromatosome | position 81 meCpA | N.D. |  |  | N.D. |  |  |
| FL MeCP2 | 15-N601-15 | H1.0<sub>1-97</sub><br>Chromatosome | position 81 meCpA | 381.7 | ± | 14.1 | 1.6 | ± | 0.1 |
| FL MeCP2 | 15-N601-15 | H1.0<sub>25-97</sub><br>Chromatosome | position 81 meCpA | 294.7 | ± | 35.7 | 1.4 | ± | 0.2 |
| FL MeCP2 AT-DIR<sub>mut</sub> | 37-N601-27 | Nucleosome | 3x meCpG | 42.5 | ± | 1.8 | 1.8 | ± | 0.1 |
| FL MeCP2 AT-DIR<sub>mut</sub> | 37-N601-27 | Nucleosome | position -1 meCpG | 119.0 | ± | 4.3 | 1.9 | ± | 0.1 |
| FL MeCP2 AT-DIR<sub>mut</sub> | 37-N601-27 | Nucleosome | unmethylated | 167.1 | ± | 5.1 | 2.1 | ± | 0.1 |
| FL MeCP2 AT-DIR<sup>mut</sup> | 15-N601-15 | Nucleosome | position -1 meCpG | 71.3 | ± | 2.3 | 3.3 | ± | 0.3 |
| FL MeCP2 AT-DIR<sup>mut</sup> | 15-N601-15 | Nucleosome | position -61 meCpG | 78.6 | ± | 2.6 | 3.1 | ± | 0.3 |
| FL MeCP2 AT-DIR<sup>mut</sup> | 15-N601-15 | Nucleosome | unmethylated | 85.2 | ± | 3.5 | 2.7 | ± | 0.3 |
| FL MeCP2 AT-DIR<sup>mut</sup> | 16-N603-30 | Nucleosome | position 91 meCpA | 25.3 | ± | 0.6 | 1.8 | ± | 0.1 |
| FL MeCP2 AT-DIR<sup>mut</sup> | 16-N603-30 | Nucleosome | position 81 meCpA | 42.2 | ± | 1.9 | 2.4 | ± | 0.2 |
| FL MeCP2 AT-DIR<sup>mut</sup> | 16-N603-30 | Nucleosome | unmethylated | 41.5 | ± | 1.4 | 2.0 | ± | 0.1 |
| MBD | 37-N601-27 | Nucleosome | 3x meCpG | 67.9 | ± | 4.6 | 1.0 | ± | 0.1 |
| MBD | 37-N601-27 | Nucleosome | position -1 meCpG | 233.7 | ± | 9.9 | 1.6 | ± | 0.1 |
| MBD | 37-N601-27 | Nucleosome | unmethylated | 278.6 | ± | 17.9 | 1.5 | ± | 0.1 |
| MBD | 15-N601-15 | Nucleosome | position -1 meCpG | 812.5 | ± | 55.7 | 1.6 | ± | 0.1 |
| MBD | 15-N601-15 | Nucleosome | position -61 meCpG | 808.9 | ± | 56.2 | 1.9 | ± | 0.2 |
| MBD | 15-N601-15 | Nucleosome | unmethylated | 1001.9 | ± | 62.9 | 1.6 | ± | 0.1 |
| MBD | N601 | Nucleosome | unmethylated | N.D. |  |  | N.D. |  |  |
| MBD | 16-N603-30 | Nucleosome | position 91 meCpA | 85.7 | ± | 6.1 | 1.0 | ± | 0.1 |
| MBD | 16-N603-30 | Nucleosome | position 81 meCpA | 267.6 | ± | 17.3 | 1.3 | ± | 0.1 |
| MBD | 16-N603-30 | Nucleosome | unmethylated | 715.9 | ± | 38.3 | 2.0 | ± | 0.2 |
| MBD | 15-N601-15 | Nucleosome | position 81 meCpA | 424.4 | ± | 14.4 | 1.5 | ± | 0.1 |
| MBD | 15-N601-15 | Nucleosome,<br>H3tailless | position 81 meCpA | 334.1 | ± | 11.7 | 1.5 | ± | 0.1 |
| MBD | 15-N601-15 | Nucleosome,<br>H3K27me3 | position 81 meCpA | 442.2 | ± | 23.7 | 1.4 | ± | 0.1 |
| FL MeCP2 R133G | 16-N603-30 | Nucleosome | position 91 meCpA | 161.6 | ± | 4.8 | 3.7 | ± | 0.3 |
| FL MeCP2 R133G | 16-N603-30 | Nucleosome | unmethylated | 187.7 | ± | 9.8 | 2.9 | ± | 0.4 |
| FL MeCP2 R133G | 15-N601-15 | Nucleosome | unmethylated | 683.0 | ± | 42.8 | 1.8 | ± | 0.2 |
| FL MeCP2 R133G | N601 | Nucleosome | unmethylated | N.D. |  |  | N.D. |  |  |
| MeCP2 162-309 | 16-N603-30 | Nucleosome | position 91 meCpA | 102.9 | ± | 2.7 | 1.9 | ± | 0.1 |
| MeCP2 162-309 | 16-N603-30 | Nucleosome | unmethylated | 94.6 | ± | 3.4 | 2.7 | ± | 0.2 |
| MeCP2 162-309 | 15-N601-15 | Nucleosome | unmethylated | 239.6 | ± | 11.8 | 2.8 | ± | 0.4 |
| MeCP2 162-309 | N601 | Nucleosome | unmethylated | 734.3 | ± | 53.4 | 2.6 | ± | 0.4 |
| MeCP2 162-309 | 15-N601-15 | Nucleosome, AP<br>mutant | unmethylated | 217.0 | ± | 9.3 | 2.4 | ± | 0.2 |
| MeCP2 162-309 | N601 | Nucleosome | unmethylated | 786.5 | ± | 28.0 | 2.5 | ± | 0.2 |
| MeCP2 162-309 | N601 | Nucleosome, AP<br>mutant | unmethylated | 1832.1 | ± | 78.3 | 3.6 | ± | 0.5 |
| MeCP2 162-309 | 40 bp | DNA | unmethylated | 31.6 | ± | 1.7 | 2.9 | ± | 0.4 |
| MeCP2 205-257 | 16-N603-30 | Nucleosome | unmethylated | 298.1 | ± | 9.5 | 2.6 | ± | 0.2 |
| MeCP2 205-257 | 15-N601-15 | Nucleosome | unmethylated | 581.2 | ± | 21.1 | 2.1 | ± | 0.1 |
| MeCP2 205-257 | N601 | Nucleosome | unmethylated | 1968.5 | ± | 229.5 | 1.3 | ± | 0.2 |
| MeCP2 205-257 | 40 bp | DNA | unmethylated | 215.5 | ± | 9.2 | 1.5 | ± | 0.1 |
| MeCP2 205-257 | 20 bp | DNA | unmethylated | 814.5 | ± | 50.1 | 1.2 | ± | 0.1 |
| MeCP2 205-257<br>K254N K256N | 40 bp | DNA | unmethylated | N.D. |  |  | N.D. |  |  |
| MeCP2 272-309 | 40 bp | DNA | unmethylated | N.D. |  |  | N.D. |  |  |
| MeCP2 258-309 | 40 bp | DNA | unmethylated | 167.3 | ± | 8.5 | 1.9 | ± | 0.2 |
| MeCP2 205-309 | 40 bp | DNA | unmethylated | 77.5 | ± | 4.8 | 2.6 | ± | 0.4 |
| DNMT3A 1-427 | 40 bp | DNA | unmethylated | 87.8 | ± | 5.2 | 2.2 | ± | 0.2 |
| DNMT3A 1-427<br>K54A K56A | 40 bp | DNA | unmethylated | 366.7 | ± | 27.5 | 1.7 | ± | 0.2 |
| HisMBP control | 16-N603-30 | Nucleosome | position 91 meCpA | N.D. |  |  | N.D. |  |  |
| HisMBP control | 16-N603-30 | Nucleosome | unmethylated | N.D. |  |  | N.D. |  |  |
| HisMBP control | 40 bp | DNA | unmethylated | N.D. |  |  | N.D. |  |  |
| HisMBP control | 20 bp | DNA | unmethylated | N.D. |  |  | N.D. |  |  |

## Notes

### Competing Interest Statement

The authors have declared no competing interest.

## References

1. Lewis, J.D. et al. Purification, sequence, and cellular localization of a novel chromosomal protein that binds to methylated DNA. Cell 69, 905–14 (1992).

2. Nan, X., Campoy, F.J. & Bird, A. MeCP2 is a transcriptional repressor with abundant binding sites in genomic chromatin. Cell 88, 471–81 (1997).

3. Nan, X. et al. Transcriptional repression by the methyl-CpG-binding protein MeCP2 involves a histone deacetylase complex. Nature 393, 386–9 (1998).

4. Lyst, M.J. et al. Rett syndrome mutations abolish the interaction of MeCP2 with the NCoR/SMRT co-repressor. Nat Neurosci 16, 898–902 (2013).

5. Kinde, B., Gabel, H.W., Gilbert, C.S., Griffith, E.C. & Greenberg, M.E. Reading the unique DNA methylation landscape of the brain: Non-CpG methylation, hydroxymethylation, and MeCP2. Proc Natl Acad Sci U S A 112, 6800–6 (2015).

6. Lagger, S. et al. MeCP2 recognizes cytosine methylated tri-nucleotide and di-nucleotide sequences to tune transcription in the mammalian brain. PLoS Genet 13, e1006793 (2017).

7. Cholewa-Waclaw, J. et al. Quantitative modelling predicts the impact of DNA methylation on RNA polymerase II traffic. Proc Natl Acad Sci U S A 116, 14995–15000 (2019).

8. Liu, Y. et al. MECP2 directly interacts with RNA polymerase II to modulate transcription in human neurons. Neuron 112, 1943–1958 e10 (2024).

9. Mishra, G.P. et al. Interaction of methyl-CpG-binding protein 2 (MeCP2) with distinct enhancers in the mouse cortex. Nat Neurosci 28, 62–71 (2025).

10. Sonn, J.Y., et al. MeCP2 Interacts with the Super Elongation Complex to Regulate Transcription. bioRxiv (2024).

11. Skene, P.J. et al. Neuronal MeCP2 is expressed at near histone-octamer levels and globally alters the chromatin state. Mol Cell 37, 457–68 (2010).

12. Kishi, N. & Macklis, J.D. MECP2 is progressively expressed in post-migratory neurons and is involved in neuronal maturation rather than cell fate decisions. Mol Cell Neurosci 27, 306–21 (2004).

13. Guy, J., Gan, J., Selfridge, J., Cobb, S. & Bird, A. Reversal of neurological defects in a mouse model of Rett syndrome. Science 315, 1143–7 (2007).

14. Fyffe, S.L. et al. Deletion of Mecp2 in Sim1-expressing neurons reveals a critical role for MeCP2 in feeding behavior, aggression, and the response to stress. Neuron 59, 947–58 (2008).

15. Ghosh, R.P., Horowitz-Scherer, R.A., Nikitina, T., Gierasch, L.M. & Woodcock, C.L. Rett syndrome-causing mutations in human MeCP2 result in diverse structural changes that impact folding and DNA interactions. J Biol Chem 283, 20523–34 (2008).

16. Guy, J. et al. A mutation-led search for novel functional domains in MeCP2. Hum Mol Genet 27, 2531–2545 (2018).

17. Amir, R.E. et al. Rett syndrome is caused by mutations in X-linked MECP2, encoding methyl-CpG-binding protein 2. Nat Genet 23, 185–8 (1999).

18. Tillotson, R. & Bird, A. The Molecular Basis of MeCP2 Function in the Brain. J Mol Biol 432, 1602–1623 (2020).

19. Guy, J., Hendrich, B., Holmes, M., Martin, J.E. & Bird, A. A mouse Mecp2-null mutation causes neurological symptoms that mimic Rett syndrome. Nat Genet 27, 322–6 (2001).

20. Akbarian, S. et al. Expression pattern of the Rett syndrome gene MeCP2 in primate prefrontal cortex. Neurobiol Dis 8, 784–91 (2001).

21. Liu, Y. et al. Exploring the complexity of MECP2 function in Rett syndrome. Nat Rev Neurosci (2025).

22. Tillotson, R. et al. Neuronal non-CG methylation is an essential target for MeCP2 function. Mol Cell 81, 1260–1275 e12 (2021).

23. Baubec, T., Ivanek, R., Lienert, F. & Schubeler, D. Methylation-dependent and −independent genomic targeting principles of the MBD protein family. Cell 153, 480–92 (2013).

24. Chen, L. et al. MeCP2 binds to non-CG methylated DNA as neurons mature, influencing transcription and the timing of onset for Rett syndrome. Proc Natl Acad Sci U S A 112, 5509–14 (2015).

25. Nan, X., Meehan, R.R. & Bird, A. Dissection of the methyl-CpG binding domain from the chromosomal protein MeCP2. Nucleic Acids Res 21, 4886–92 (1993).

26. Ho, K.L. et al. MeCP2 binding to DNA depends upon hydration at methyl-CpG. Mol Cell 29, 525–31 (2008).

27. Sperlazza, M.J., Bilinovich, S.M., Sinanan, L.M., Javier, F.R. & Williams, D.C., Jr. Structural Basis of MeCP2 Distribution on Non-CpG Methylated and Hydroxymethylated DNA. J Mol Biol 429, 1581–1594 (2017).

28. Lei, M., Tempel, W., Chen, S., Liu, K. & Min, J. Plasticity at the DNA recognition site of the MeCP2 mCG-binding domain. Biochim Biophys Acta Gene Regul Mech 1862, 194409 (2019).

29. Xie, W. et al. Base-resolution analyses of sequence and parent-of-origin dependent DNA methylation in the mouse genome. Cell 148, 816–31 (2012).

30. Varley, K.E. et al. Dynamic DNA methylation across diverse human cell lines and tissues. Genome Res 23, 555–67 (2013).

31. Guo, H. et al. The DNA methylation landscape of human early embryos. Nature 511, 606–10 (2014).

32. Ballestar, E., Yusufzai, T.M. & Wolffe, A.P. Effects of Rett syndrome mutations of the methyl-CpG binding domain of the transcriptional repressor MeCP2 on selectivity for association with methylated DNA. Biochemistry 39, 7100–6 (2000).

33. Fraga, M.F. et al. The affinity of different MBD proteins for a specific methylated locus depends on their intrinsic binding properties. Nucleic Acids Res 31, 1765–74 (2003).

34. Ibrahim, A. et al. MeCP2 is a microsatellite binding protein that protects CA repeats from nucleosome invasion. Science 372(2021).

35. Ghosh, R.P. et al. Unique physical properties and interactions of the domains of methylated DNA binding protein 2. Biochemistry 49, 4395–410 (2010).

36. Adams, V.H., McBryant, S.J., Wade, P.A., Woodcock, C.L. & Hansen, J.C. Intrinsic disorder and autonomous domain function in the multifunctional nuclear protein, MeCP2. J Biol Chem 282, 15057–64 (2007).

37. Baker, S.A. et al. An AT-hook domain in MeCP2 determines the clinical course of Rett syndrome and related disorders. Cell 152, 984–96 (2013).

38. Heckman, L.D., Chahrour, M.H. & Zoghbi, H.Y. Rett-causing mutations reveal two domains critical for MeCP2 function and for toxicity in MECP2 duplication syndrome mice. Elife 3(2014).

39. Lyst, M.J., Connelly, J., Merusi, C. & Bird, A. Sequence-specific DNA binding by AT-hook motifs in MeCP2. FEBS Lett 590, 2927–33 (2016).

40. Claveria-Gimeno, R. et al. The intervening domain from MeCP2 enhances the DNA affinity of the methyl binding domain and provides an independent DNA interaction site. Sci Rep 7, 41635 (2017).

41. Ortega-Alarcon, D. et al. Influence of the disordered domain structure of MeCP2 on its structural stability and dsDNA interaction. Int J Biol Macromol 175, 58–66 (2021).

42. McGinty, R.K. & Tan, S. Nucleosome structure and function. Chem Rev 115, 2255–73 (2015).

43. Chandler, S.P., Guschin, D., Landsberger, N. & Wolffe, A.P. The methyl-CpG binding transcriptional repressor MeCP2 stably associates with nucleosomal DNA. Biochemistry 38, 7008–18 (1999).

44. Ishibashi, T., Thambirajah, A.A. & Ausio, J. MeCP2 preferentially binds to methylated linker DNA in the absence of the terminal tail of histone H3 and independently of histone acetylation. FEBS Lett 582, 1157–62 (2008).

45. Nikitina, T. et al. MeCP2-chromatin interactions include the formation of chromatosome-like structures and are altered in mutations causing Rett syndrome. J Biol Chem 282, 28237–45 (2007).

46. Georgel, P.T. et al. Chromatin compaction by human MeCP2. Assembly of novel secondary chromatin structures in the absence of DNA methylation. J Biol Chem 278, 32181–8 (2003).

47. Yang, C., van der Woerd, M.J., Muthurajan, U.M., Hansen, J.C. & Luger, K. Biophysical analysis and small-angle X-ray scattering-derived structures of MeCP2-nucleosome complexes. Nucleic Acids Res 39, 4122–35 (2011).

48. Chua, G.N.L. et al. Differential dynamics specify MeCP2 function at nucleosomes and methylated DNA. Nat Struct Mol Biol 31, 1789–1797 (2024).

49. Bartke, T. et al. Nucleosome-interacting proteins regulated by DNA and histone methylation. Cell 143, 470–84 (2010).

50. Thambirajah, A.A. et al. MeCP2 binds to nucleosome free (linker DNA) regions and to H3K9/H3K27 methylated nucleosomes in the brain. Nucleic Acids Res 40, 2884–97 (2012).

51. Lee, W., Kim, J., Yun, J.M., Ohn, T. & Gong, Q. MeCP2 regulates gene expression through recognition of H3K27me3. Nat Commun 11, 3140 (2020).

52. Ortega-Alarcon, D. et al. Extending MeCP2 interactome: canonical nucleosomal histones interact with MeCP2. Nucleic Acids Res 52, 3636–3653 (2024).

53. Tillotson, R. et al. Radically truncated MeCP2 rescues Rett syndrome-like neurological defects. Nature 550, 398–401 (2017).

54. Lowary, P.T. & Widom, J. New DNA sequence rules for high affinity binding to histone octamer and sequence-directed nucleosome positioning. J Mol Biol 276, 19–42 (1998).

55. Kan, P.Y., Lu, X., Hansen, J.C. & Hayes, J.J. The H3 tail domain participates in multiple interactions during folding and self-association of nucleosome arrays. Mol Cell Biol 27, 2084–91 (2007).

56. Peng, Y., Li, S., Onufriev, A., Landsman, D. & Panchenko, A.R. Binding of regulatory proteins to nucleosomes is modulated by dynamic histone tails. Nat Commun 12, 5280 (2021).

57. Piccolo, F.M. et al. MeCP2 nuclear dynamics in live neurons results from low and high affinity chromatin interactions. Elife 8(2019).

58. McGinty, R.K. & Tan, S. Principles of nucleosome recognition by chromatin factors and enzymes. Curr Opin Struct Biol 71, 16–26 (2021).

59. Kumar, A. et al. Analysis of protein domains and Rett syndrome mutations indicate that multiple regions influence chromatin-binding dynamics of the chromatin-associated protein MECP2 in vivo. J Cell Sci 121, 1128–37 (2008).

60. Klose, R.J. et al. DNA binding selectivity of MeCP2 due to a requirement for A/T sequences adjacent to methyl-CpG. Mol Cell 19, 667–78 (2005).

61. Agarwal, N. et al. MeCP2 Rett mutations affect large scale chromatin organization. Hum Mol Genet 20, 4187–95 (2011).

62. Pantier, R. et al. MeCP2 binds to methylated DNA independently of phase separation and heterochromatin organisation. Nat Commun 15, 3880 (2024).

63. Landrum, M.J. et al. ClinVar: public archive of relationships among sequence variation and human phenotype. Nucleic Acids Res 42, D980–5 (2014).

64. Ben Chorin, A., et al. ConSurf-DB: An accessible repository for the evolutionary conservation patterns of the majority of PDB proteins. Protein Sci 29, 258–267 (2020).

65. Cheng, J. et al. Accurate proteome-wide missense variant effect prediction with AlphaMissense. Science 381, eadg7492 (2023).

66. Tordai, H. et al. Analysis of AlphaMissense data in different protein groups and structural context. Sci Data 11, 495 (2024).

67. Zhou, X. et al. A novel mutation R190H in the AT-hook 1 domain of MeCP2 identified in an atypical Rett syndrome. Oncotarget 8, 82156–82164 (2017).

68. Wang, T. et al. De novo genic mutations among a Chinese autism spectrum disorder cohort. Nat Commun 7, 13316 (2016).

69. Suetake, I. et al. Characterization of DNA-binding activity in the N-terminal domain of the DNA methyltransferase Dnmt3a. Biochem J 437, 141–8 (2011).

70. Turlure, F., Maertens, G., Rahman, S., Cherepanov, P. & Engelman, A. A tripartite DNA-binding element, comprised of the nuclear localization signal and two AT-hook motifs, mediates the association of LEDGF/p75 with chromatin in vivo. Nucleic Acids Res 34, 1653–65 (2006).

71. Wapenaar, H. et al. The N-terminal region of DNMT3A combines multiple chromatin reading motifs to guide recruitment. bioRxiv, 2023.10.29.564595 (2023).

72. Wapenaar, H. et al. The N-terminal region of DNMT3A engages the nucleosome surface to aid chromatin recruitment. EMBO Rep 25, 5743–5779 (2024).

73. Wang, H., Farnung, L., Dienemann, C. & Cramer, P. Structure of H3K36-methylated nucleosome-PWWP complex reveals multivalent cross-gyre binding. Nat Struct Mol Biol 27, 8–13 (2020).

74. Ito-Ishida, A. et al. MeCP2 Levels Regulate the 3D Structure of Heterochromatic Foci in Mouse Neurons. J Neurosci 40, 8746–8766 (2020).

75. Pearson, E.C., Bates, D.L., Prospero, T.D. & Thomas, J.O. Neuronal nuclei and glial nuclei from mammalian cerebral cortex. Nucleosome repeat lengths, DNA contents and H1 contents. Eur J Biochem 144, 353–60 (1984).

76. Dominguez, V., Pina, B. & Suau, P. Histone H1 subtype synthesis in neurons and neuroblasts. Development 115, 181–5 (1992).

77. Ghosh, R.P., Horowitz-Scherer, R.A., Nikitina, T., Shlyakhtenko, L.S. & Woodcock, C.L. MeCP2 binds cooperatively to its substrate and competes with histone H1 for chromatin binding sites. Mol Cell Biol 30, 4656–70 (2010).

78. Lister, R. et al. Global epigenomic reconfiguration during mammalian brain development. Science 341, 1237905 (2013).

79. Riedmann, C. & Fondufe-Mittendorf, Y.N. Comparative analysis of linker histone H1, MeCP2, and HMGD1 on nucleosome stability and target site accessibility. Sci Rep 6, 33186 (2016).

80. Chereji, R.V., Ramachandran, S., Bryson, T.D. & Henikoff, S. Precise genome-wide mapping of single nucleosomes and linkers in vivo. Genome Biol 19, 19 (2018).

81. Clark, S.C., Chereji, R.V., Lee, P.R., Fields, R.D. & Clark, D.J. Differential nucleosome spacing in neurons and glia. Neurosci Lett 714, 134559 (2020).

82. Baldi, S., Korber, P. & Becker, P.B. Beads on a string-nucleosome array arrangements and folding of the chromatin fiber. Nat Struct Mol Biol 27, 109–118 (2020).

83. Lyst, M.J. et al. Affinity for DNA Contributes to NLS Independent Nuclear Localization of MeCP2. Cell Rep 24, 2213–2220 (2018).

84. Burdett, H. et al. BRCA1-BARD1 combines multiple chromatin recognition modules to bridge nascent nucleosomes. Nucleic Acids Res 51, 11080–11103 (2023).

85. Dyer, P.N. et al. Reconstitution of nucleosome core particles from recombinant histones and DNA. Methods Enzymol 375, 23–44 (2004).

86. Wilson, M.D. et al. Retroviral integration into nucleosomes through DNA looping and sliding along the histone octamer. Nat Commun 10, 4189 (2019).

87. Zhou, B.R. et al. Distinct Structures and Dynamics of Chromatosomes with Different Human Linker Histone Isoforms. Mol Cell 81, 166–182 e6 (2021).

88. Wilson, M.D. et al. The structural basis of modified nucleosome recognition by 53BP1. Nature 536, 100–3 (2016).

89. Deak, G. et al. Histone divergence in trypanosomes results in unique alterations to nucleosome structure. Nucleic Acids Res (2023).

90. Turner, T. Plot protein: visualization of mutations. J Clin Bioinforma 3, 14 (2013).

91. Deak, G. & Cook, A.G. Missense Variants Reveal Functional Insights Into the Human ARID Family of Gene Regulators. J Mol Biol 434, 167529 (2022).

92. Sigrist, C.J. et al. New and continuing developments at PROSITE. Nucleic Acids Res 41, D344–7 (2013).

93. Altschul, S.F. et al. Gapped BLAST and PSI-BLAST: a new generation of protein database search programs. Nucleic Acids Res 25, 3389–402 (1997).

94. Rappsilber, J., Mann, M. & Ishihama, Y. Protocol for micro-purification, enrichment, pre-fractionation and storage of peptides for proteomics using StageTips. Nat Protoc 2, 1896–906 (2007).

95. Chambers, M.C. et al. A cross-platform toolkit for mass spectrometry and proteomics. Nat Biotechnol 30, 918–20 (2012).

96. Mendes, M.L. et al. An integrated workflow for crosslinking mass spectrometry. Mol Syst Biol 15, e8994 (2019).

97. Fischer, L. & Rappsilber, J. Quirks of Error Estimation in Cross-Linking/Mass Spectrometry. Anal Chem 89, 3829–3833 (2017).

98. Vasudevan, D., Chua, E.Y.D. & Davey, C.A. Crystal structures of nucleosome core particles containing the ‘601’ strong positioning sequence. J Mol Biol 403, 1–10 (2010).

99. Klose, R.J. & Bird, A.P. MeCP2 behaves as an elongated monomer that does not stably associate with the Sin3a chromatin remodeling complex. J Biol Chem 279, 46490–6 (2004).

100. Connelly, J.C. et al. Absence of MeCP2 binding to non-methylated GT-rich sequences in vivo. Nucleic Acids Res 48, 3542–3552 (2020).

101. Schmiedeberg, L., Skene, P., Deaton, A. & Bird, A. A temporal threshold for formaldehyde crosslinking and fixation. PLoS One 4, e4636 (2009).

102. Belotserkovskaya, R. et al. PALB2 chromatin recruitment restores homologous recombination in BRCA1-deficient cells depleted of 53BP1. Nat Commun 11, 819 (2020).

