## Supplementary figures and images for "MeCP2 requires interactions with nucleosome linker DNA to read chromatin DNA methylation"

### Extended Data Figures

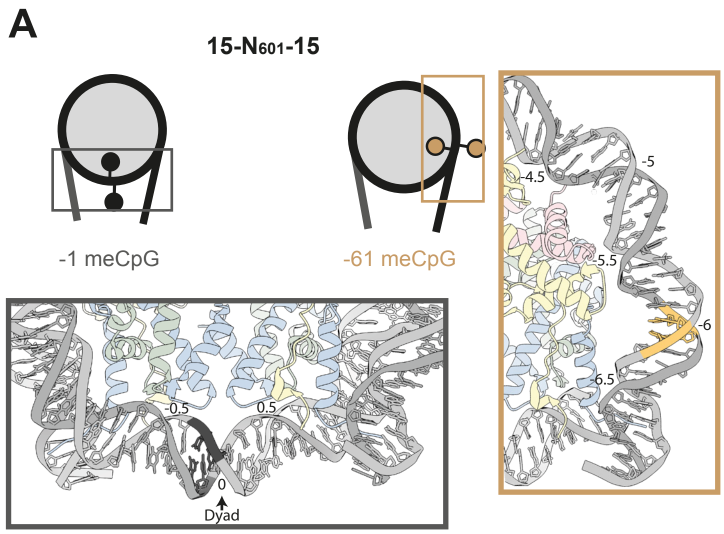

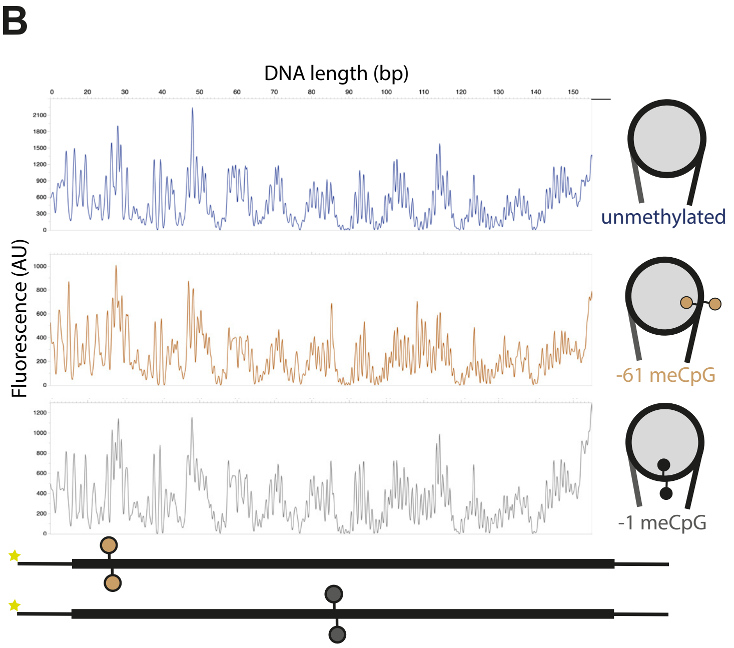

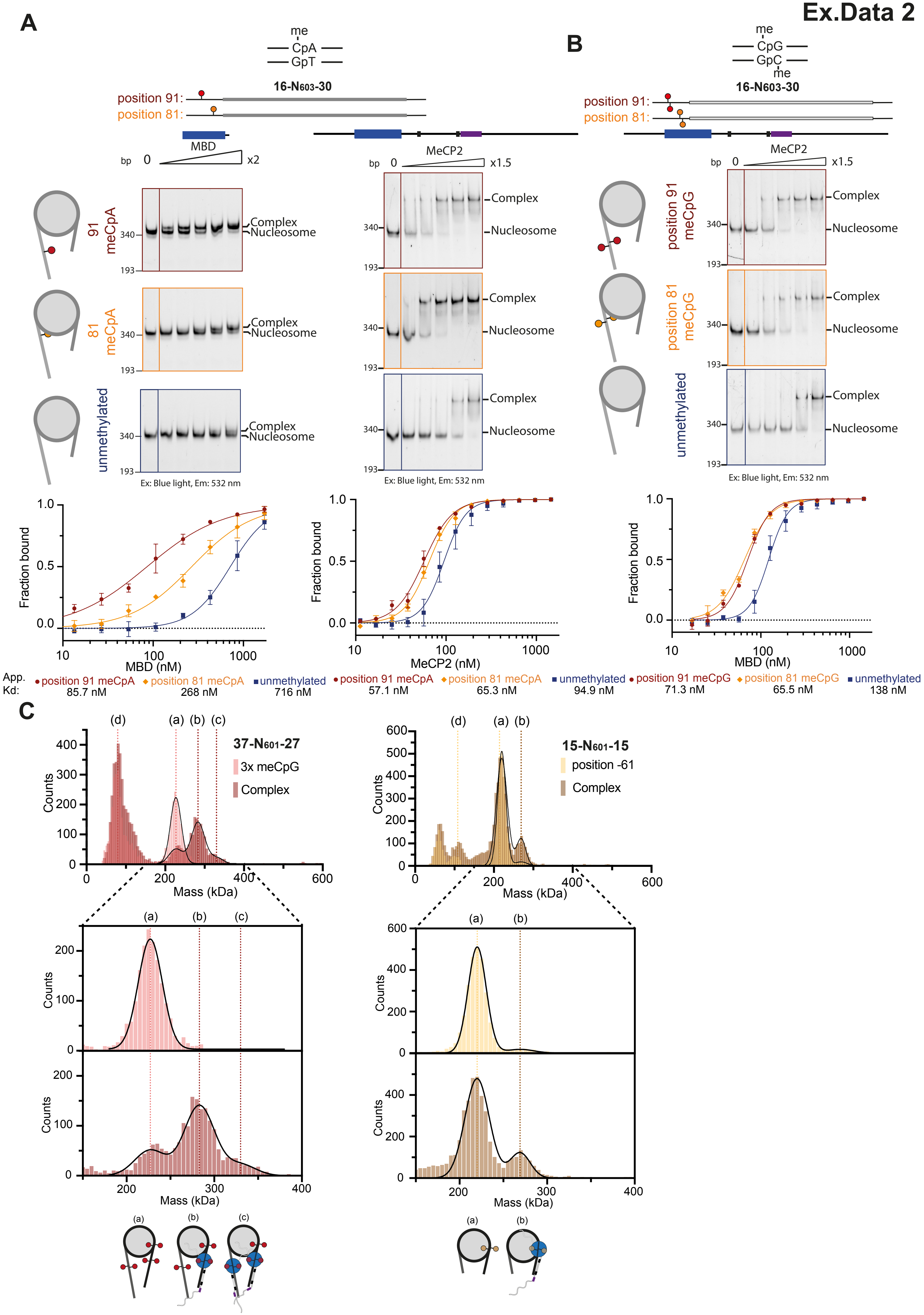

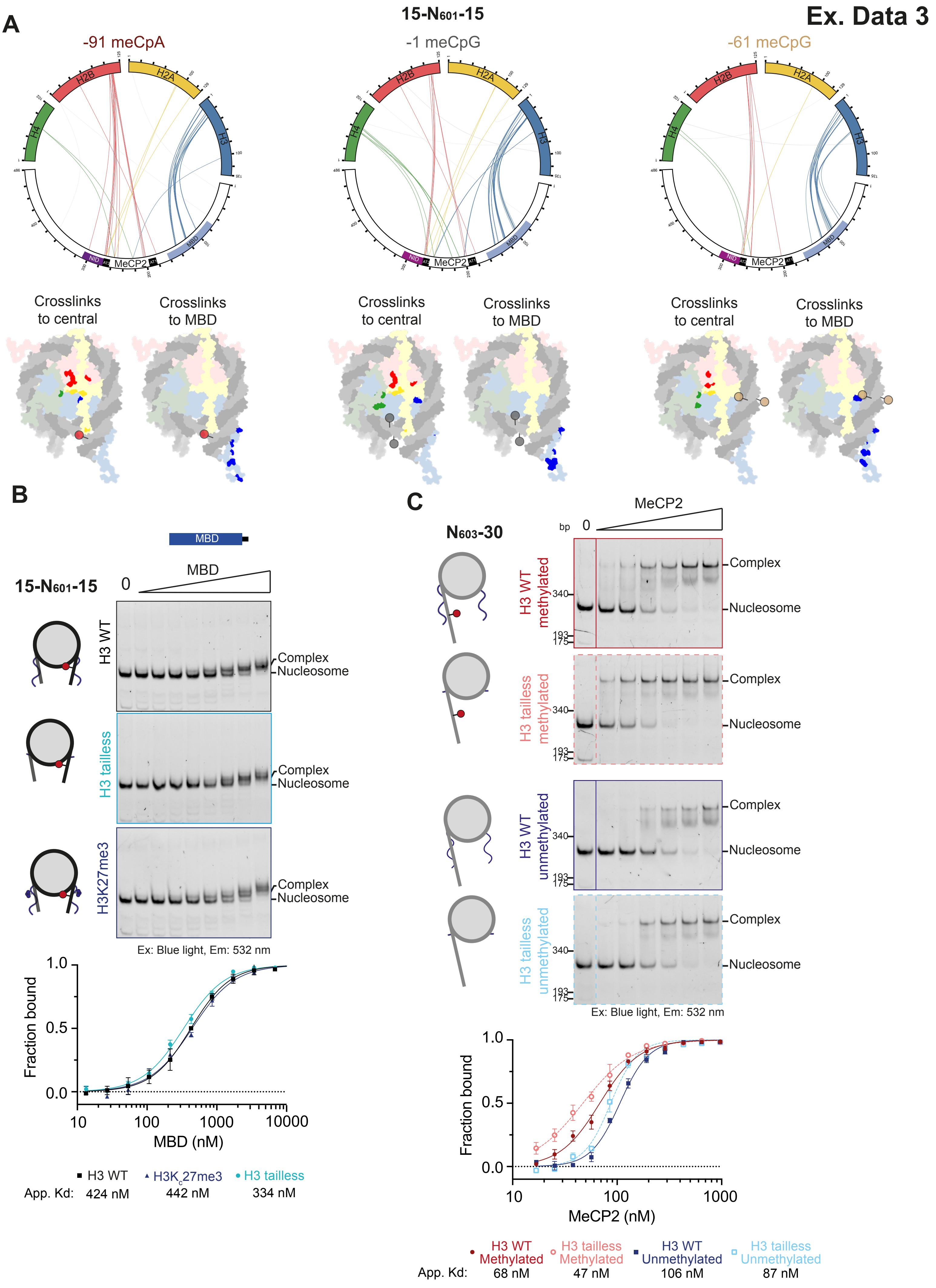

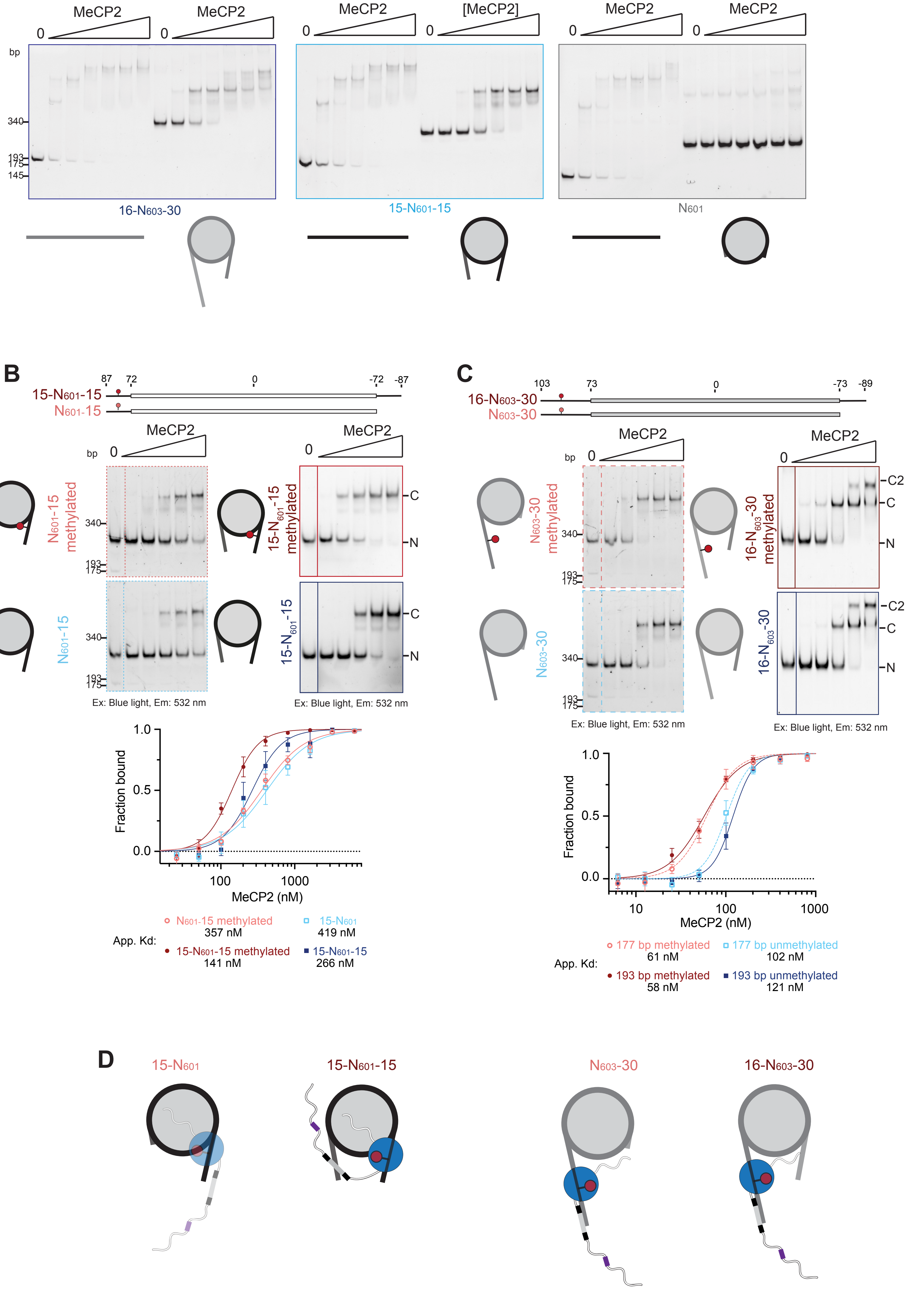

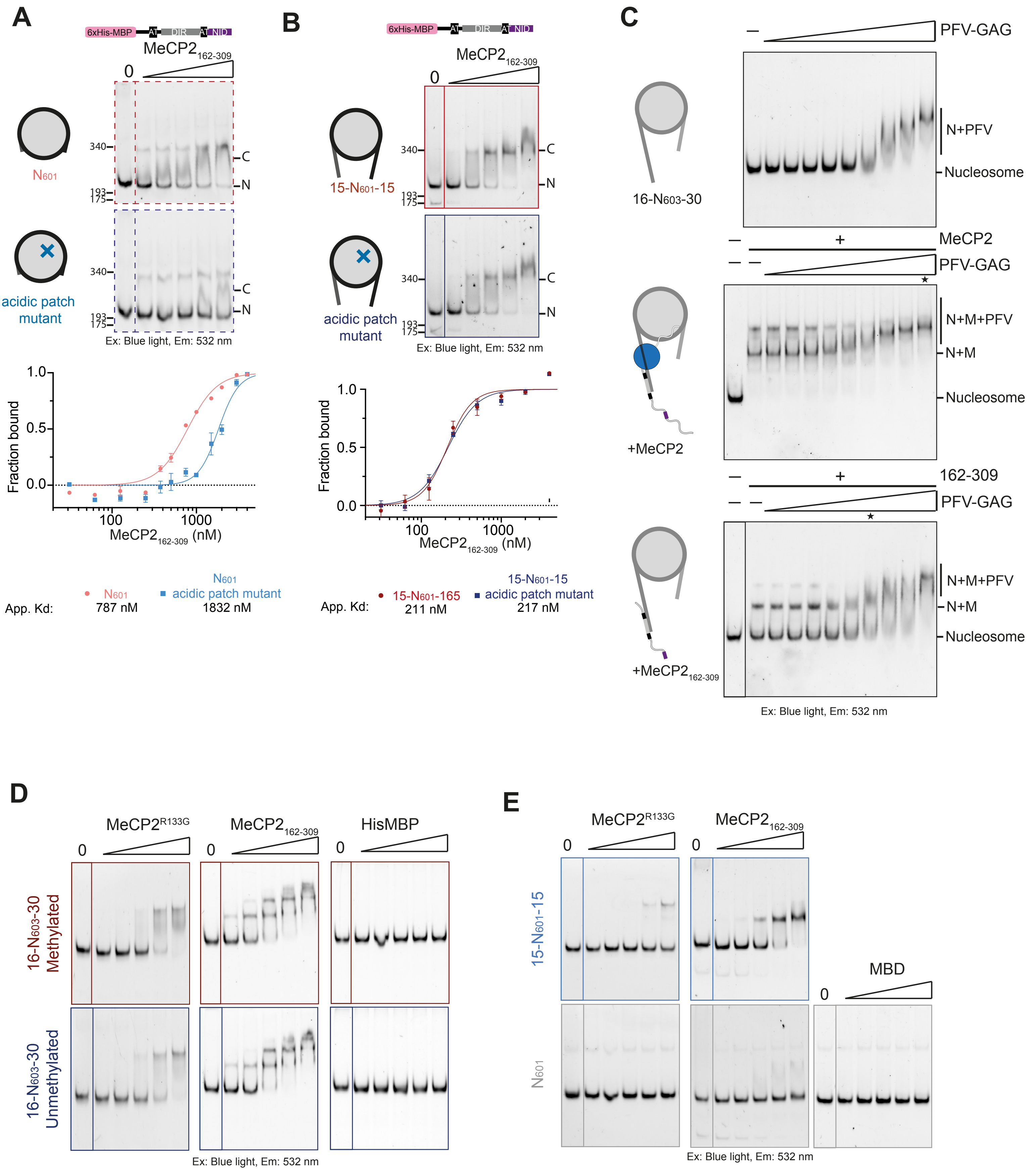

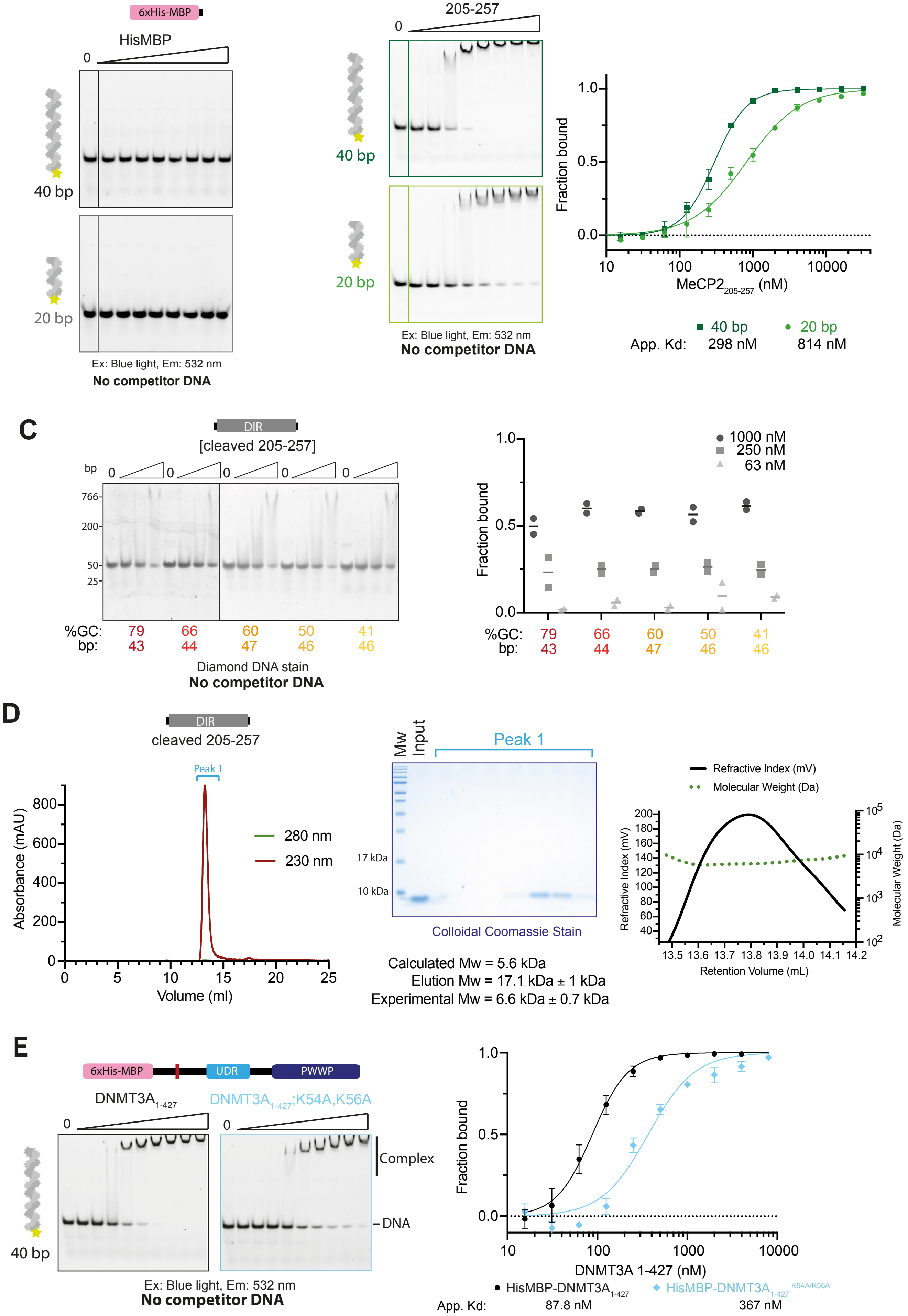

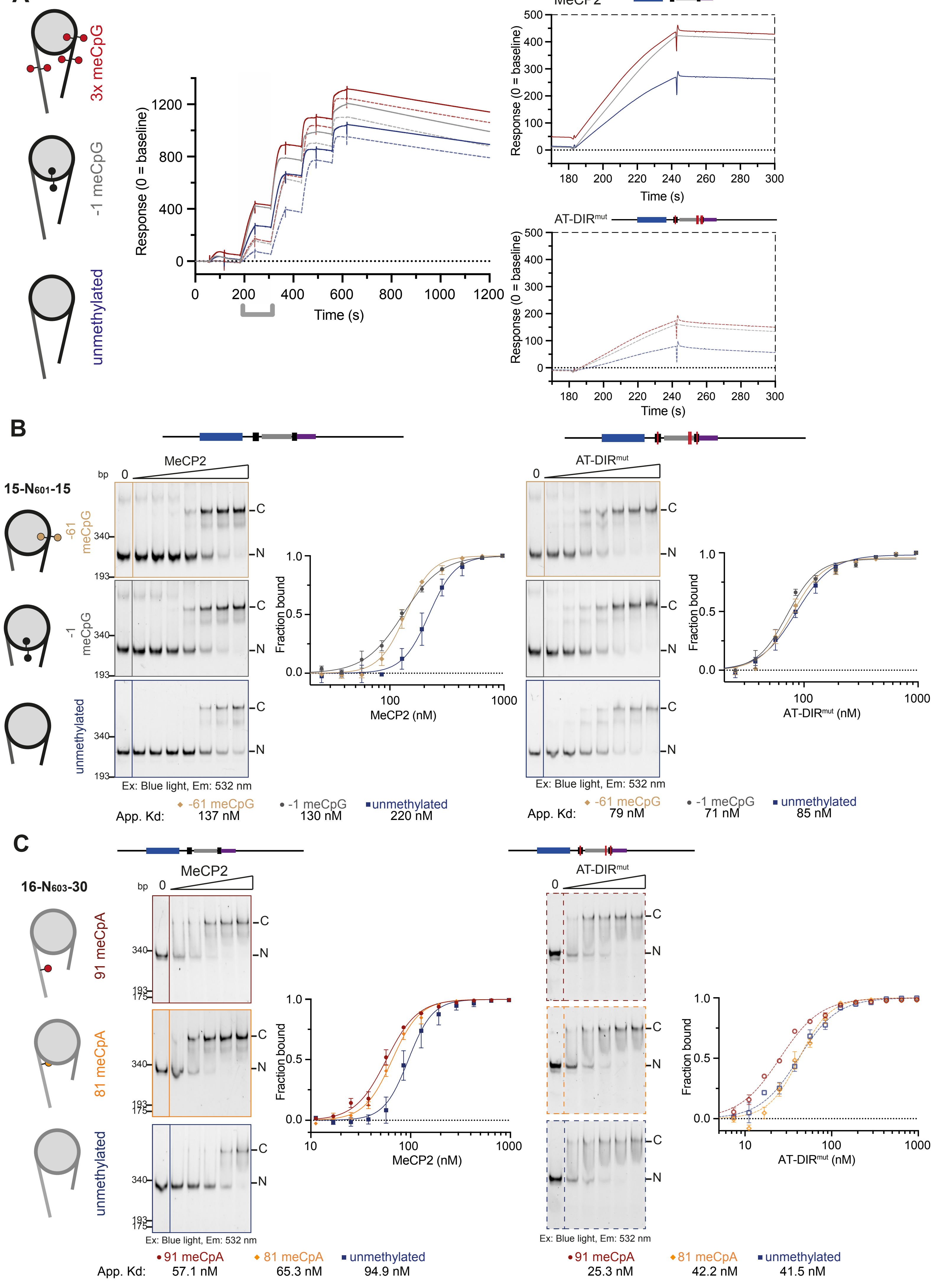

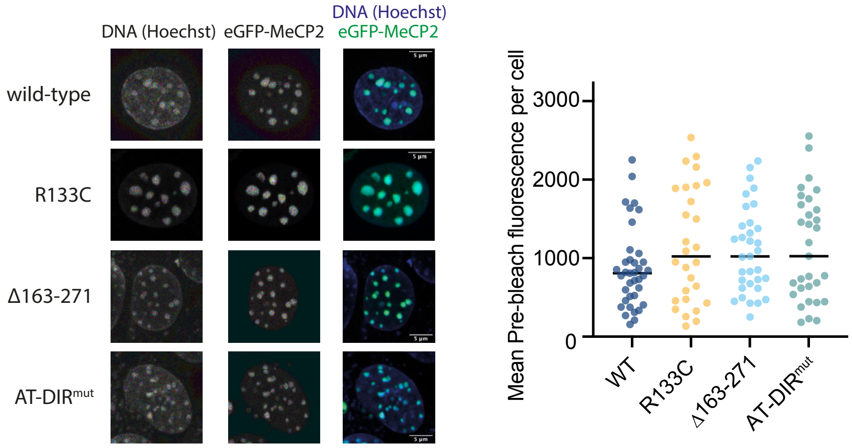

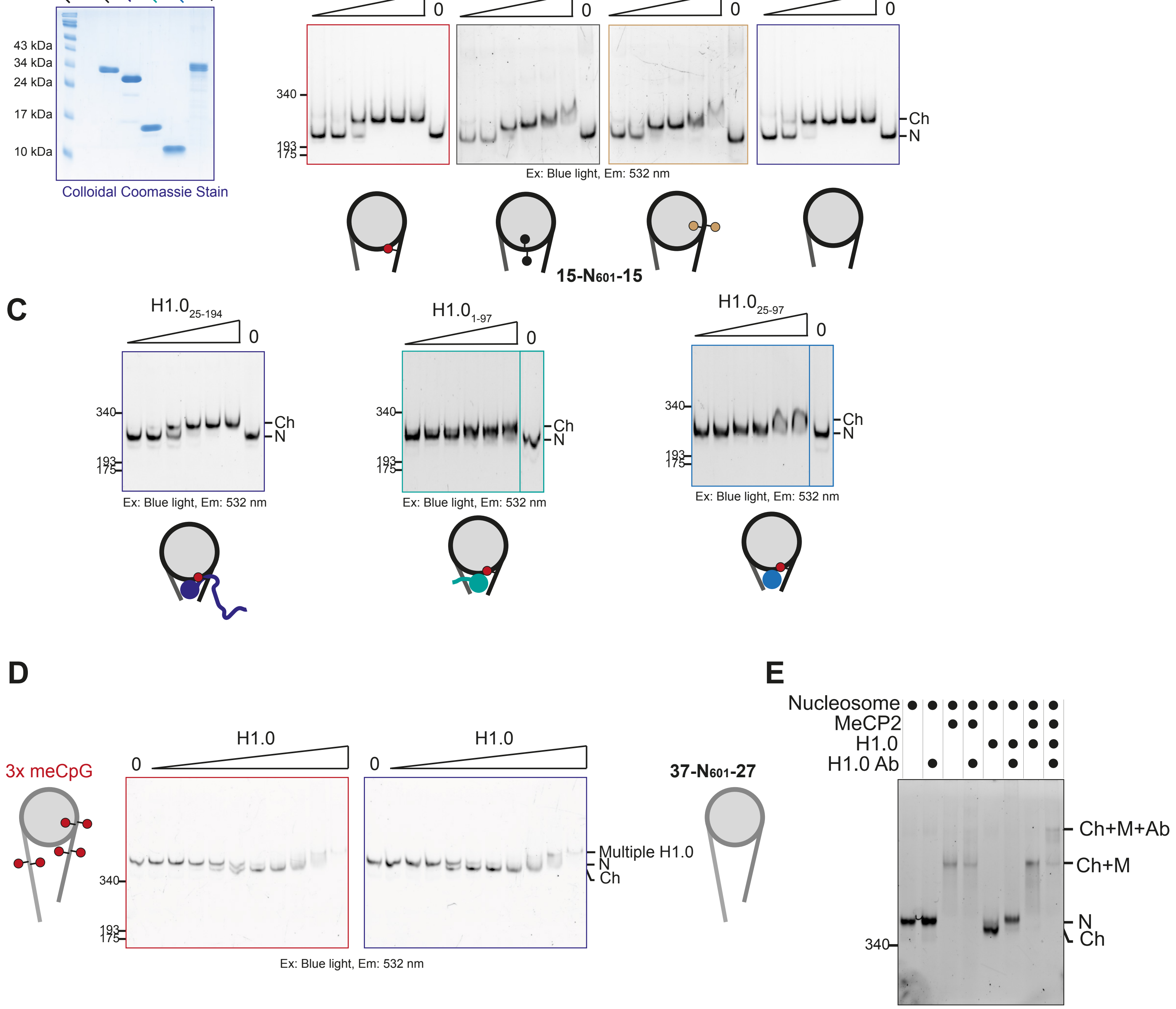

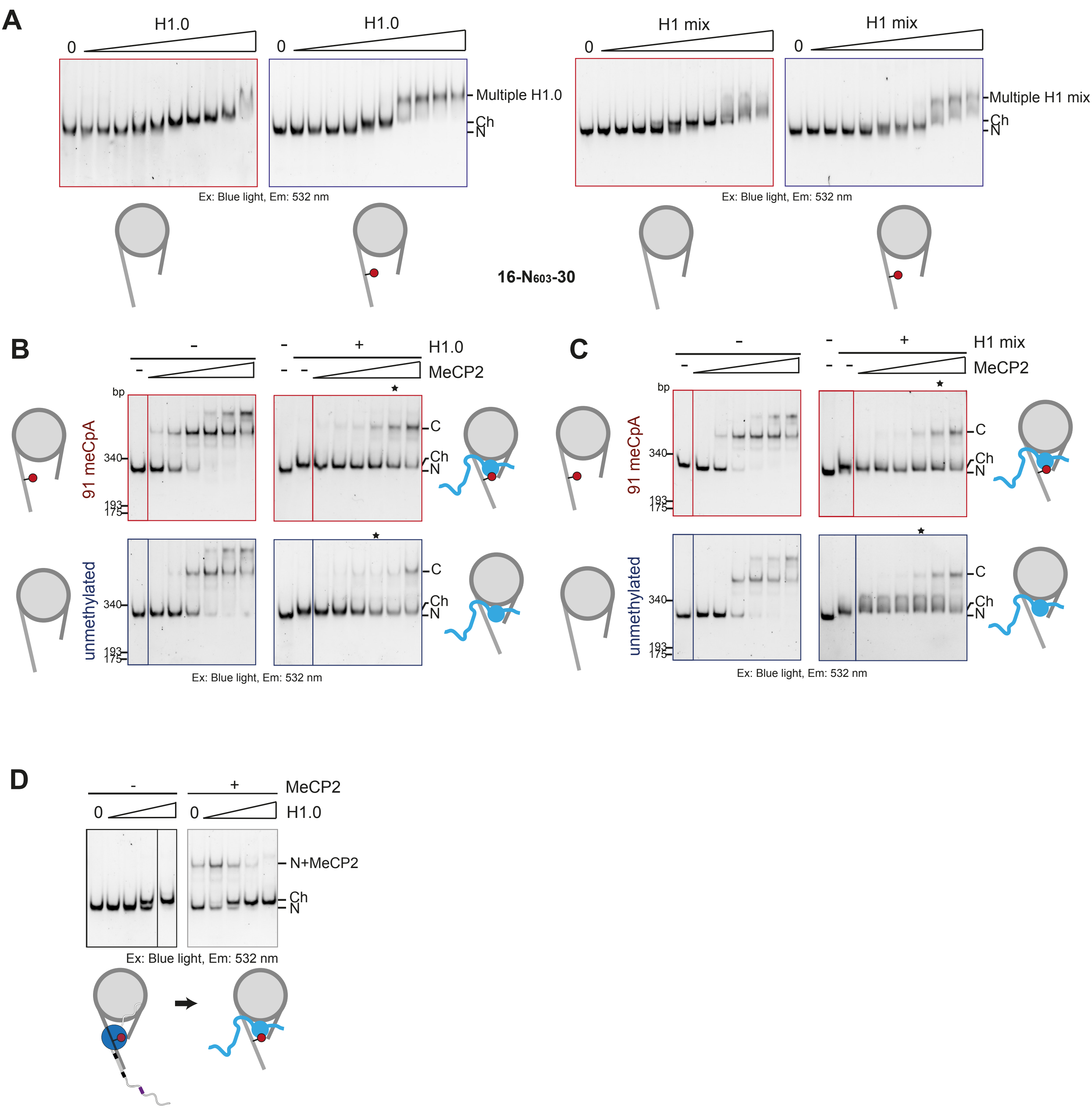

### Supplemental Figures

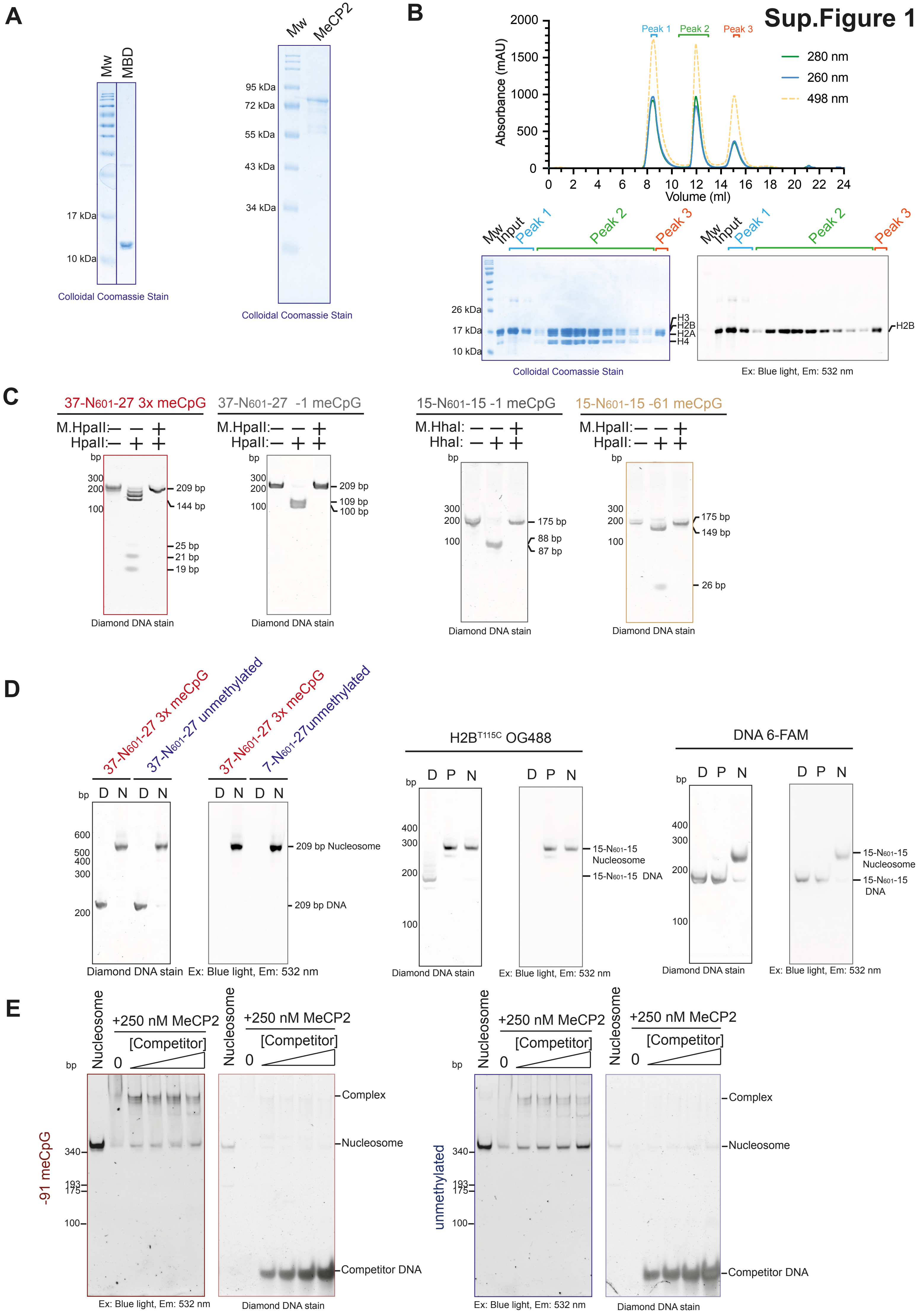

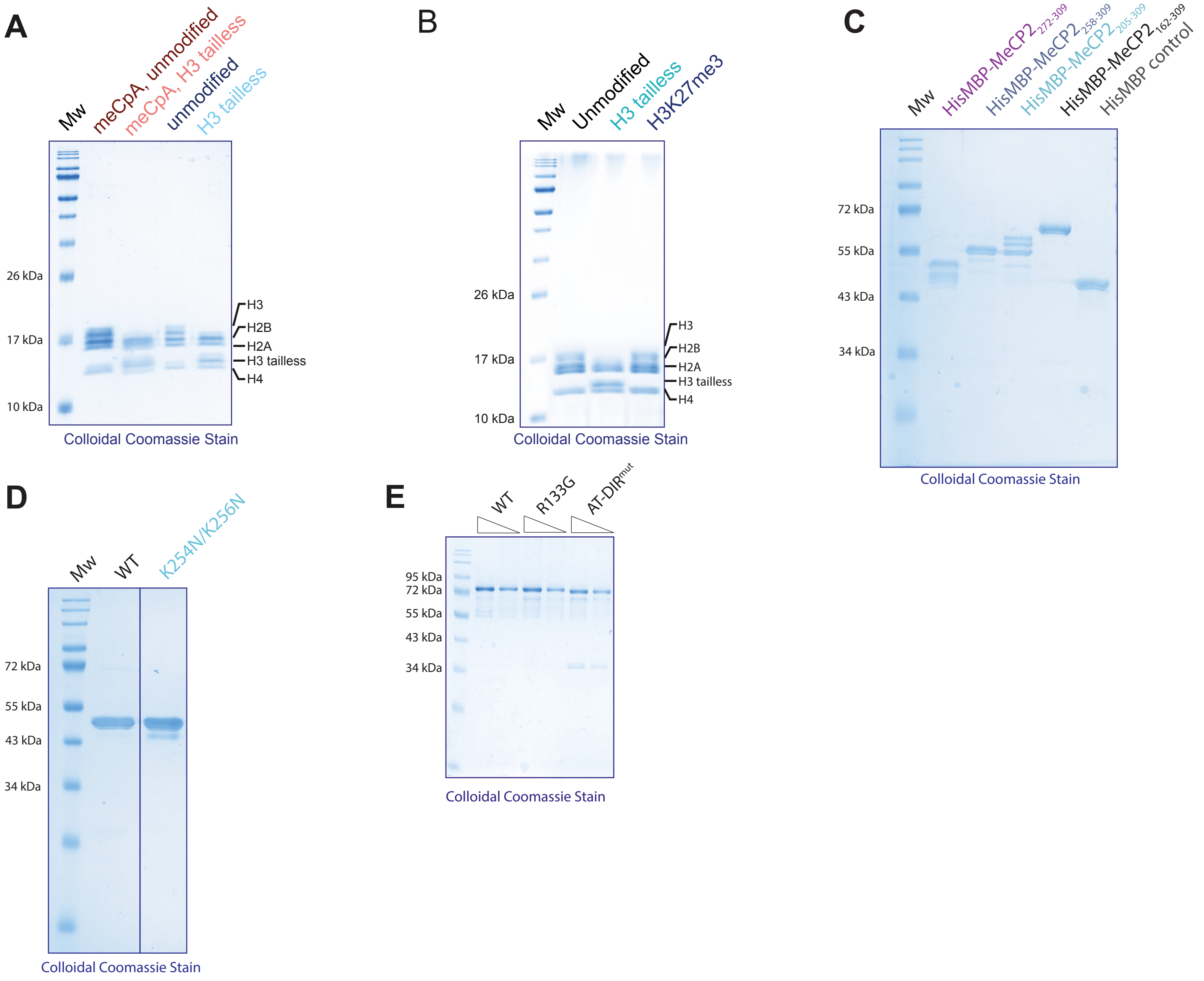

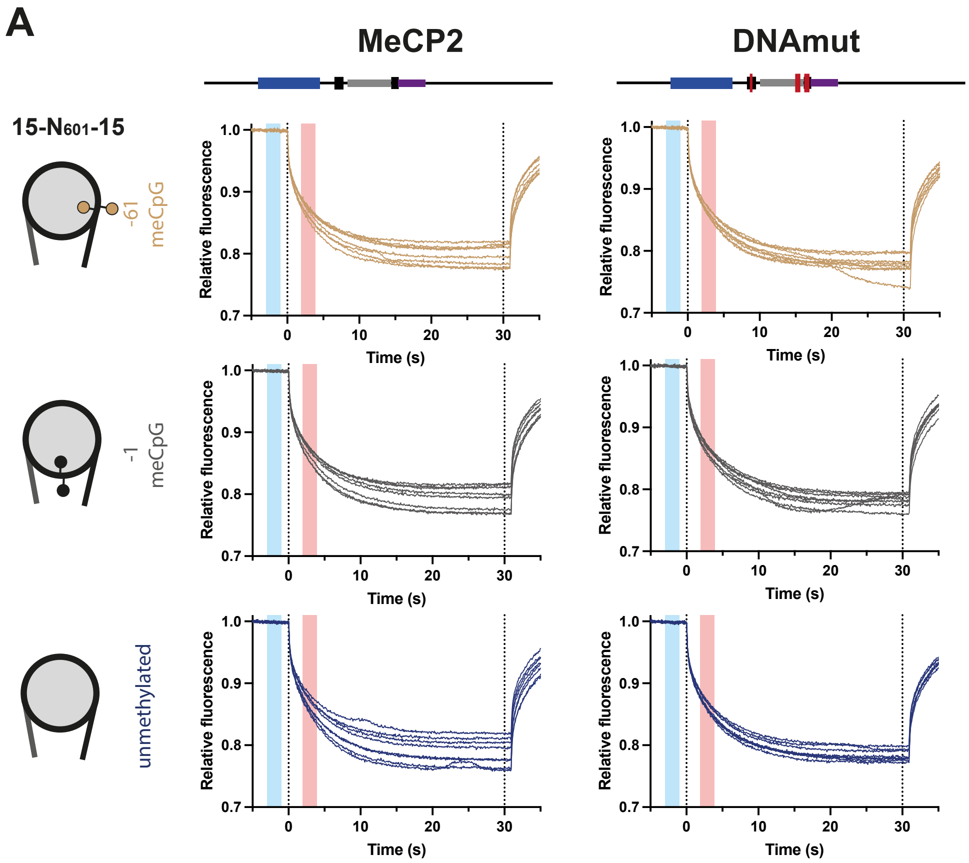
